# Psychophysical measurement of perceived motion flow of naturalistic scenes

**DOI:** 10.1101/2023.02.14.528582

**Authors:** Yung-Hao Yang, Taiki Fukiage, Zitang Sun, Shin’ya Nishida

## Abstract

The neural and computational mechanisms underlying visual motion perception have been extensively investigated over several decades, but most studies have used simple artificial stimuli such as random-dot kinematograms. Thus, it remains difficult to predict how human observers perceive optical flows in complex natural scenes. Here, we report a novel method to measure, psychophysically, optical flows perceived by human observers watching naturalistic movies, and to reveal the characteristics of human motion perception via comparison of the measured perceived flow to the ground truths and model predictions. We selected movie clips from the MPI Sintel Flow Dataset, which contains open-source computer graphics animations with ground truths. To measure the perceived vectors at a spatiotemporal point, we flashed a small dot during presentation of a brief clip and asked the observers to adjust the speed and direction of a matching random-noise stimulus, to reproduce the vector at the flashed point. The proposed method adequately estimated perceived flow, and the estimated perceived vector also indicated flow illusions, i.e., consistent deviations from the ground truths, in various ways, depending on the stimulus patterns. Comparisons with the predictions of biologically motivated models and machine vision algorithms indicated that some flow illusions were attributable to lower-level factors such as spatiotemporal pooling and signal loss, but others reflected higher-level computations including coordinate transformations that cannot be precisely predicted by existing flow estimation models. Psychophysical measurement of the optical flows that humans perceive in realistic environments constitutes a promising paradigm for advancing our understanding of visual motion perception.

**Significance Statement:** The basic approach to studying human vision is to analyze relationships among subjective perceptual experiences, responses of neural mechanisms, and predictions of computational models. Recent technical advances have enabled researchers to access large-scale neuronal responses and model predictions for complex sensory inputs. The data on human visual perception, however, cannot be easily scaled up. Accurate measurement of rich visual experiences when viewing natural scenes remains challenging. We thus devised a novel psychophysical method to measure optical motion flows perceived by humans. We successfully visualized human-perceived flows of complex naturalistic movies and show, for the first time, ways in which human-perceived naturalistic flows agree with, and deviate from, the physical ground truths and the predictions of various visual motion models.

## Introduction

Visual motion perception is one of the most extensively investigated cognitive functions (see refs. 1–4 for reviews). Past studies have revealed that the first stage of visual motion processing is extraction of directional signals by spatiotemporally oriented sensors (5), followed by mutual integration and inhibition of such signals, and estimation of an array of image motion vectors that is also termed an optical flow map (6). The higher-level visual motion processing uses this map to recognize object motion in the scene, to control eye and body movements, to self-navigate in the field via optical flow, and to perceive biological motion. The cortical mechanisms underlying these computations have been also extensively studied (7). Progress in visual motion research has been aided by the use of several artificial stimuli that selectively tap each processing stage; the stimuli include drifting sine-wave gratings (8), plaids (9), random-dot kinematograms (10–12), point-light walkers (13), and global dot flow patterns (14).

Recent technical advances have rendered it possible to access large-scale data on neural responses, and to create models that predict the responses to complex stimuli. Interest in visual motion research has thus shifted to how the visual system processes complex visual motion information in dynamic natural environments. For instance, Nishimoto & Gallant (15) measured the cortical functional magnetic resonance imaging (fMRI) responses to movie clips of natural scenes, and tested the validity of the models having been proposed to explain human motion perception. Matthis et al. (16) measured visual motions projected on the retina during natural locomotion when studying the role of optical flow in action control in real-world environments.

However, to the best of our knowledge, few attempts have been made to measure psychophysically optical flows that human observers really perceive in complex natural or naturalistic scenes. Such perceived flows, if measurable, would advance our understanding of visual motion perception. It would be possible to compare human perceptions directly with the ground truths (GTs), neurophysiologically measured brain responses, and the predictions of the several image-computable motion models developed in the field of vision science and machine vision.

Psychophysical estimation of a perceived optical flow map is challenging because this requires measurements of perceived vectors (speed and direction) at many spatiotemporal positions in a dynamic scene. Here, we present a novel measurement procedure, inspired by the gauge-probe task of surface shape estimation (17). Observers are asked to report perceived vectors by adjusting the speed and direction of a matching noise stimulus. To indicate the target spatiotemporal position during each trial, we superimpose a brief dot probe on the movie stimulus. By changing the target position in a grid-like fashion, we derive a map of the perceived optical flow. Objective evaluation of human performance requires the GT, but it is not easy to obtain physically correct optical flow maps for ordinal natural movies. We therefore used a synthesized, naturalistic movie dataset (18) that has been widely used for training and evaluation of optical flow models in the field of computer vision.

We found that, using this procedure, human observers reproduced the speed and direction of a motion vector at a specified spatiotemporal location with reasonable accuracy. Of interest, we found several “flow illusions” where the estimated perceived flow deviated systematically from the GT. We explored how accurately the illusions were predicted via spatiotemporal pooling of local motion signals, by visual motion models motivated by biological computations, and by computer vision models engineered for optical flow estimation. We found that some “illusions” were predicted by retinotopic, optical flow estimation models, but others reflected high-level computations including coordinate transformation and vector decomposition. Our data demonstrate the strengths and limitations of existing models that predict visual motion flow perceptions when viewing naturalistic scenes, and that estimation of human-perceived optical flow maps of a variety of natural scenes will advance our understanding of human visual motion processing.

## Results

In the main experiment, we measured the human perceived optical flows in five movie clips selected from a high-frame-rate version of the MPI Sintel Flow Dataset (18, 42). The clips feature naturalistic video sequences with multiple objects made of various materials performing different movements (Fig. 1C). The Dataset includes GT optical flow data (Fig. 1D) that can be directly compared to human responses.

**Fig. 1.**
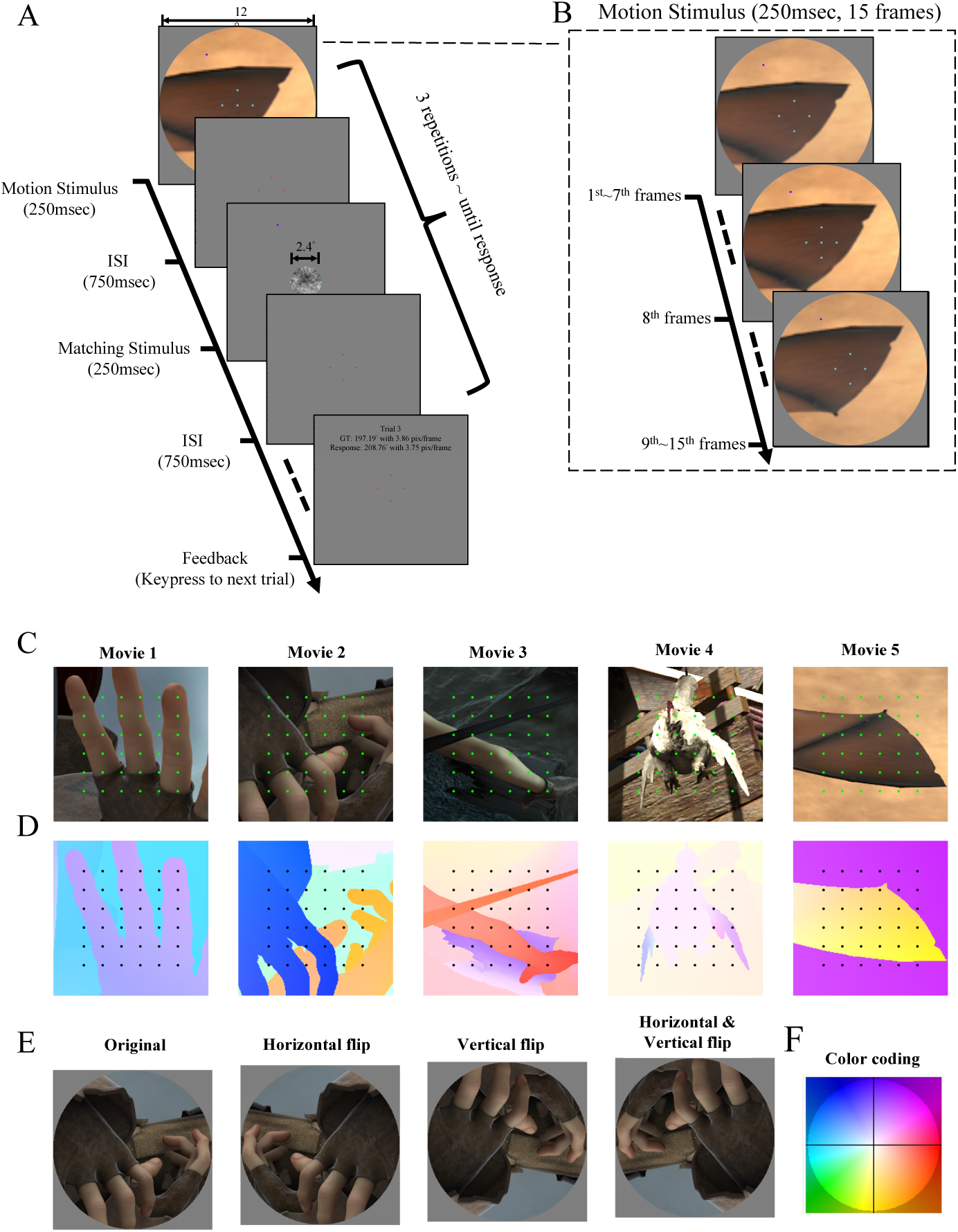
Experimental procedures and visual stimuli. (A) In each trial, motion and matching stimuli were alternatively and repeatedly presented. Each repetition featured a motion stimulus (250 ms), an inter-stimulus interval (ISI, 750 ms), a matching stimulus (250 ms), and an ISI (750 ms). (B) A probe dot was flashed in the middle (frame 8) of the motion stimulus presentation (250 ms, 15 frames) and observers were required to report the motion vector at the location/time of the flash by adjusting the speed and direction of the matching stimulus. After at least three repetitions, the observers could terminate the trial when satisfied with their settings. A feedback display was then presented. (C) The 36 probed locations (1° spacing, green dots) of each movie clip. (D) GT motion vectors of each movie using a color-coding scheme (ref. 43) where hue indicates the motion direction and saturation the speed (normalized by the maximum speed of each five clips) as shown in (F). (E) Each movie clip was presented through a circular aperture (12° in diameter) in one of four flip modes: Original orientation, horizontal flip, vertical flip, and horizontal and vertical flip.

The human-perceived optical flow map was measured by a local vector-matching method (Fig. 1 and Movie S1). In each trial, pairs of target motion clips and matching noise stimuli were repeatedly presented. The observers were asked to adjust the direction and speed of the matching stimulus to reproduce the local motion vector in the target movie clip at the location indicated by a flashed probe (Fig. 1A, B). The probe location was selected from a 6 x 6 grid placed on the region of interest (Figs 1C, D). Each movie clip was presented in one of four flip modes (Fig. 1E); this allowed us to decompose the human response errors into two components. One component of errors is defined on the display coordinates, and determined by the relationship between the observer and the display. The other component is defined on the image coordinates, the directions of which flip as the image flips. Our principal interest was the latter type of human error.

### Accuracy evaluation

We evaluated the accuracy of our novel method from two perspectives (see SI 1 and Figs. S1-S4 for further details). First, to evaluate how accurately the observers reproduced the direction and speed of the perceived flow (vector-matching accuracy), we analyzed the data of a preliminary experiment using a random-dot kinematogram. The results indicated that the reported motions agreed well with the GT motions (Fig. 2a). Next, to evaluate how accurately in space and time the observers reported local vectors specified by the flashed probe (flash-probing accuracy), we analyzed the data of the main experiment using the MPI Sintel movie clips. The results indicated that the reported motions best agreed with the GT local motions at spatiotemporal locations very close to the probe (Fig. 2c). Thus, our method yields reasonably accurate estimates of the perceived flow we attempted to measure.

**Fig. 2.**
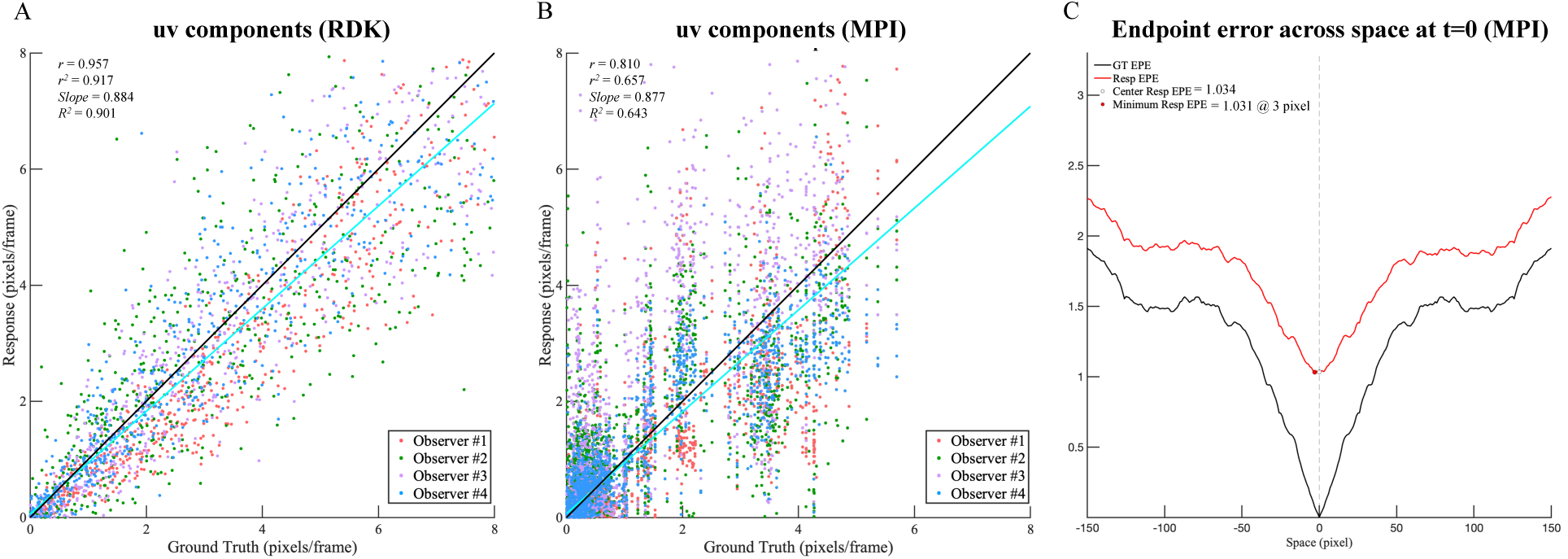
Evaluation of two aspects of accuracy. See also SI 1. (A) Vector-matching accuracy as shown by a scatter plot of responses versus the GTs for the horizontal and vertical (*uv*) vector components based on the results of a preliminary experiment using a random-dot kinematogram. Each point indicates a response of one observer in one trial. Different colors refer to different observers. *r, r^2^*: Pearson correlations and the squares; slope: the slope of the linear regression line; R2: the coefficient of determination by reference to the ideal response (the GT). The index values were computed based on the signed *uv* values; the Figure plots the absolute (unsigned) *uv* values for visualization. See the separate plots of the direction and speed components in Figure S1. (B) A similar scatter plot of the results of the main experiment using the MPI Sintel dataset. Vector-matching accuracy is reduced by flow illusions. See also Figure S2. (C) Flash-probing accuracy as shown by the spatiotemporal distribution of the endpoint differences in the display coordinates of the MPI Sintel Flow Dataset. The red line indicates the endpoint differences between the human response to a probe and the GTs at surrounding locations. The observer responses were closest to the GTs at the probed locations, indicating that observers reported motions at probed locations rather accurately. The black line indicates the within-stimulus similarity, thus the endpoint differences between the GT at the probed location and the GTs at surrounding locations. See also Figures S3 and S4.

### Human perceived flow: An overview

Figure 3 shows the response flow map for each MPI Sintel movie clip. We averaged the response vectors over the four flip conditions after unflipping them into the original image coordinates to reveal the response biases associated with the stimulus pattern while controlling for the biases associated with the relationship between the observer and the display. The results of the four observers were averaged. The pattern of results was generally similar across the observers (see SI 2, Fig. S5, and Table S1).

**Fig. 3.**
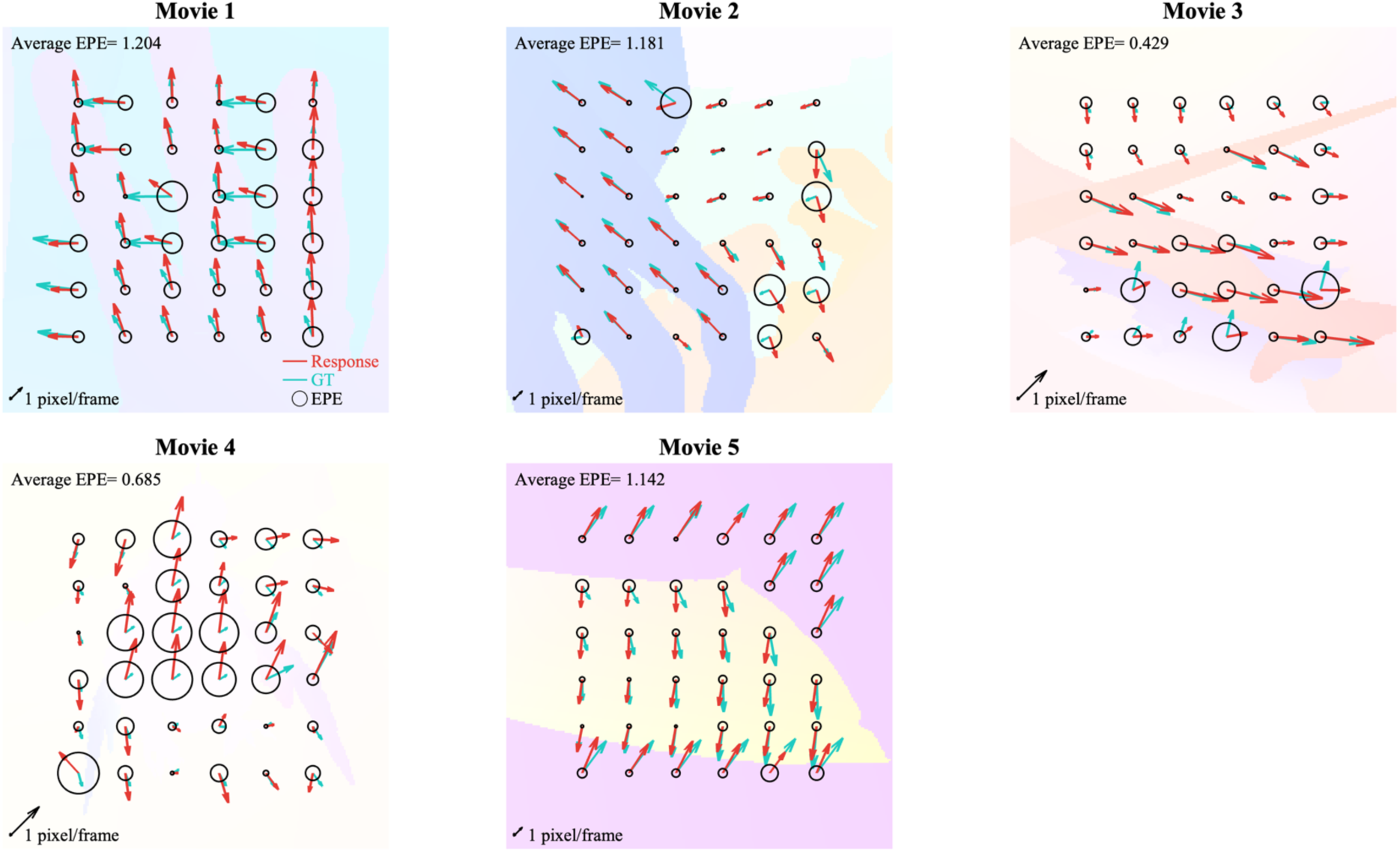
The perceived vector map for the MPI Sintel Flow Dataset. The response flow map using the image coordinates of the MPI Sintel Flow Dataset. At each location, the green arrow denotes the GT, the red arrow the human response, and the diameter of the black circle the EPE. In each panel, the average EPE of the 36 movie locations is shown in the top left corner. As the vector length was normalized within each movie, the spatial scale is shown by an arrow (of length 1 pixel/frame) in the bottom left corner.

In Figure 3, the response flow vector (red arrow) agrees with the GT vector (green arrow) at certain probed locations. The Endpoint Error (EPE) averaged over all probe points is 0.928 pixels or 1.11 arcmin (see the first line of Table 1), suggesting that the observers reliably reproduced certain flow patterns in naturalistic scenes. However, the reported vector deviated from the GT vectors at many other locations in various ways. For example, large errors are found in the gaps between fingers in Movie 1, in the hand on the right in Movie 2, in the dark background surrounding the arm in Movie 3, in the body of the rooster in Movie 4, and in the wing of the bat in Movie 5. Thus, the agreement between the reported and GT vectors was less in the main MPI SINTEL experiment than in the preliminary RDK experiment (compare Fig. 2a and 2b).

**Table 1.**
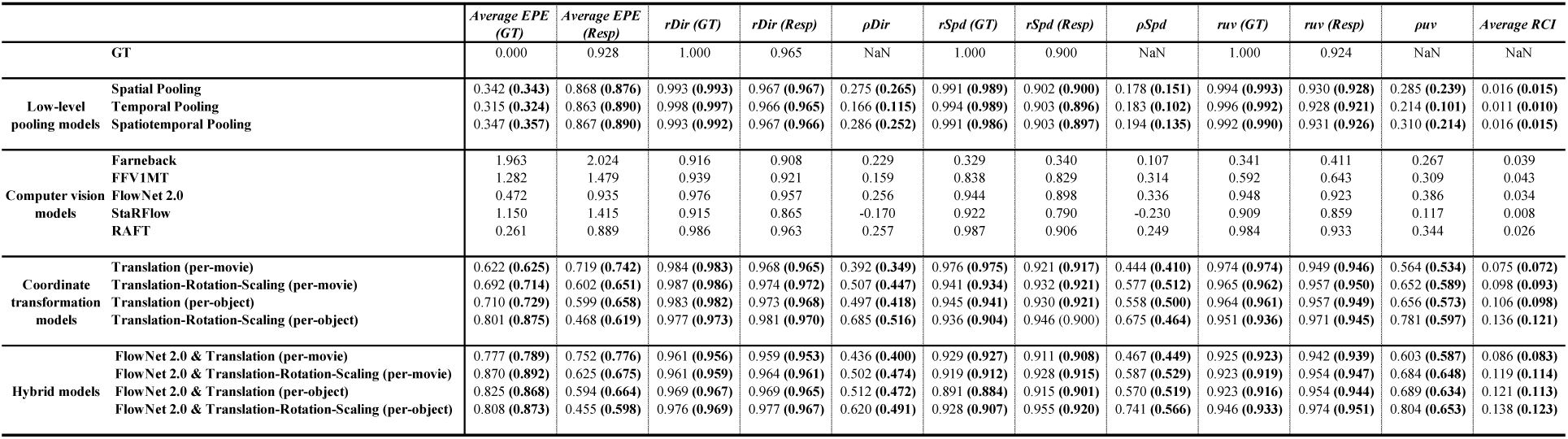
Predictive performances of the models. EPE (GT): EPE between the model prediction and the GT. EPE (Resp): EPE between the model prediction and the human response. r_Dir/Spd/uv_(GT): Pearson correlations of the direction (Dir), speed (Spd), and uv components (uv) between the model predictions and GTs. r_Dir/Spd/uv_(Resp): Pearson correlations of direction, speed, and the uv components between the model predictions and the human responses. ρ_Dir/Spd/uv_: Partial correlations of the direction, speed, and uv components between the model predictions and the human responses with the effects of GT removed. RCI: Response Consistency Index. For the fitting functions, in addition to values estimated for models fitted to all data, the medians of the estimated values computed via two-fold cross-validation are shown in bold within brackets.

### How to compare human perceived flow and model predictions?

The patterns of systematic deviation between the human vector and GT vectors, which we term “flow illusions”, would be expected to provide valuable information on human visual motion processing. To seek the mechanisms underlying the illusions, we compared the human response to the predictions of visual processing models including pooling models that simulate biologically plausible lower-level visual processing, computer vision models developed for optical flow estimation, and coordinate transformation models that simulate biologically plausible higher-level visual processing.

As shown in Table 1, to evaluate the performance of each model, we used predicted vectors to compute: (1) the average EPE by reference to the GTs averaged over all 180 probed locations; (2) the average EPE by reference to the human responses; (3) the Pearson correlations (*r* values) between the GT direction, speed, and *uv* components; (4) the Pearson correlations between the human-perceived direction, speed, and *uv* components; (5) the partial correlations (*ρ*) between the human-perceived direction, speed, and *uv* components with the effects of GT removed; and, (6) the average response consistency index (RCI) over the probed locations. With the exception of the computer vision models, we estimated the free model parameters that best explained the response data (see Table S2 for the best-fit parameters for each model). To ensure fair comparisons across models with different numbers of free parameters, we also computed index values via two-fold cross-validation, and used (principally) these values for model evaluation.

Of the indices shown in Table 1, two are most informative in terms of the extent of overall agreement between the model predictions and human responses. One is the partial correlation of *uv* components between the model predictions and human responses (*ρ_uv_*). Although the Pearson correlation between the model predictions and human responses (*r_uv_*) could be high simply because both correlate with the common variable GT, *ρ_uv_* evaluates the correlation between the two while excluding the effect of GT.

The other informative index is the spatially averaged RCI. The RCI is an index of consistency between model predictions and the human responses, and takes a value between -1 and +1. The RCI becomes positive and approaches +1 as the prediction becomes closer to the human response than to the GT, approaches zero when the model prediction becomes closer to the GT, and becomes negative when the model predicts an error in the opposite direction (see the Materials and Methods). We found a strong positive correlation between the average RCI and the partial correlation (Fig. 4). An advantage of the RCI compared to the partial correlation is that the RCI can be separately computed for each location. In addition, our analysis suggests that the average RCI is less affected by the small number of outliers than is the partial correlation.

**Fig. 4.**
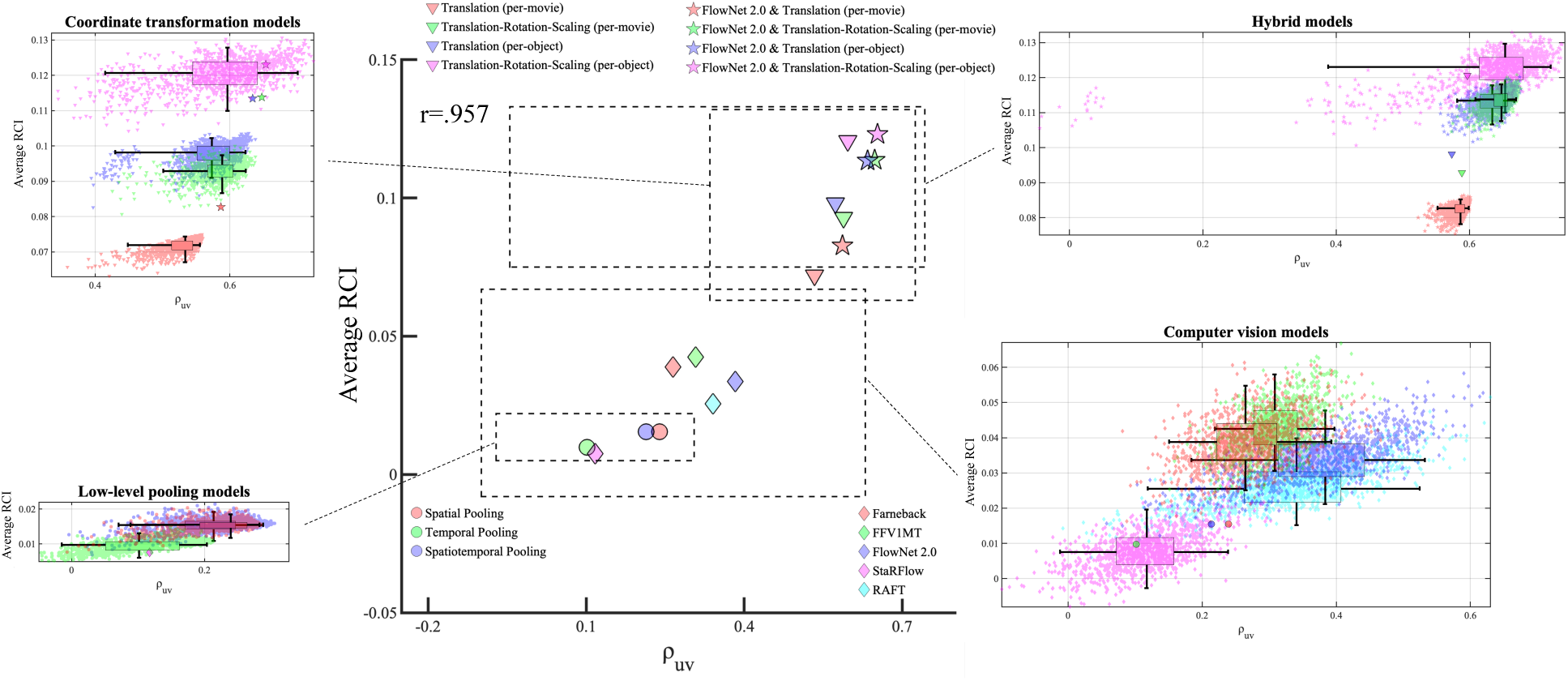
Scatter plots of the partial correlations of the *uv* components (ρ_uv_) versus the average RCIs. In the center panel, each point represents the median performance index distribution computed by the bootstrapping method for the computer vision models and by the two-fold cross-validation for the fitting models. The other four panels show the performance index distributions of 1,000 sampling sets and 2D box plots with the [25% percentile, median, 75% percentile] ranges (boxes) and the 95% confidence intervals (whiskers).

Figure 4 shows the distributions of the partial correlation in *uv* components (*ρ_uv_*) and the spatially averaged RCI, computed using the bootstrapping method (for computer vision models) and via two-fold cross-validation (for the other models). See Figure S6 for the distribution of each index. During statistical testing of the differences in index values, we checked whether zero was included in the 95% confidence intervals of the differences between each model pair computed by the bootstrapping method (for computer models) or the cross-validation method (other models) (Fig. S7).

Using the RCIs, we visualized how well the models explained the human response errors at each movie probe location (see Fig. 5 for the representative four models, and Fig. S8 for all models). To compare the similarities of model predictions, we computed the correlations of the spatial RCI patterns between each model pair (Table S3).

**Fig. 5.**
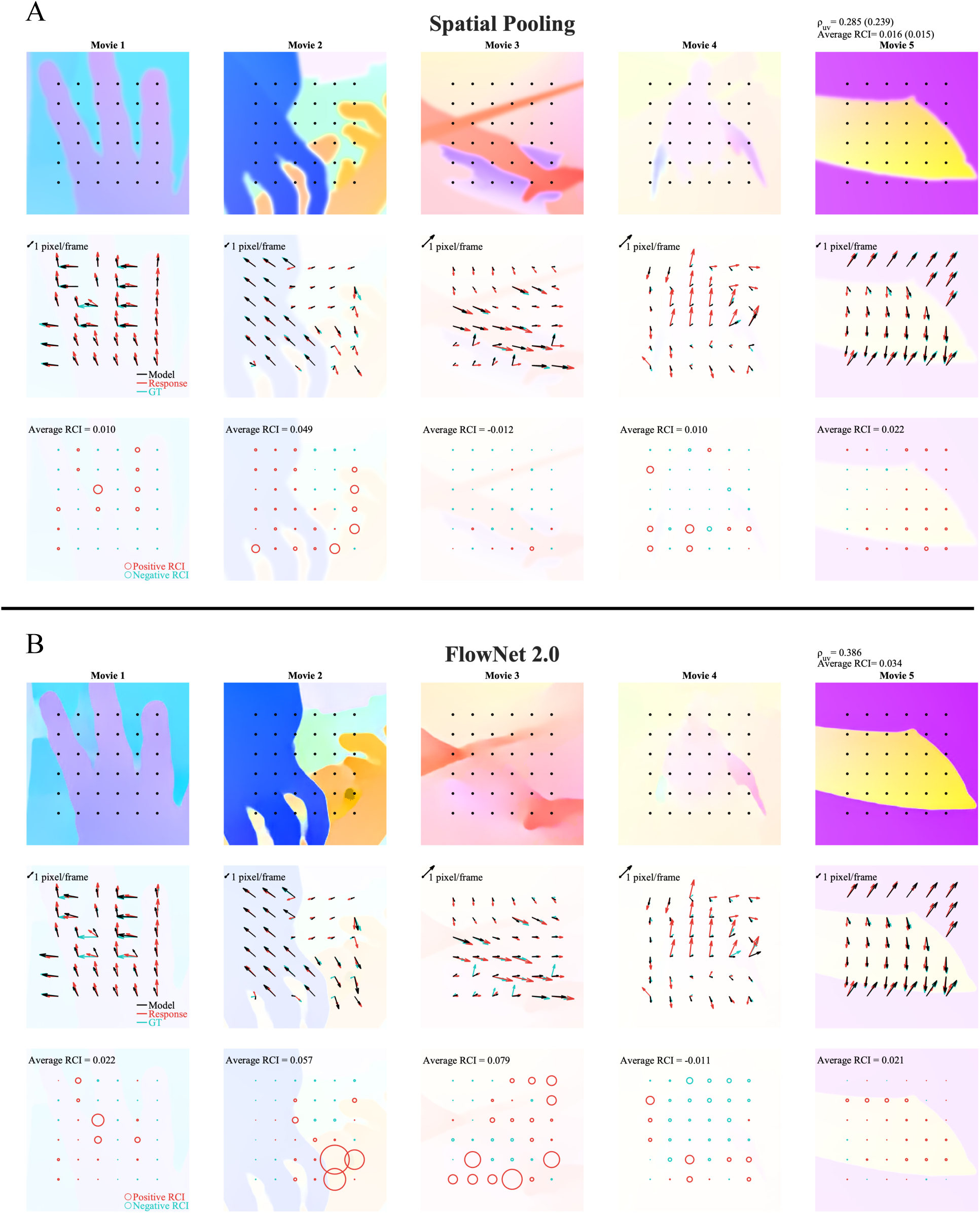
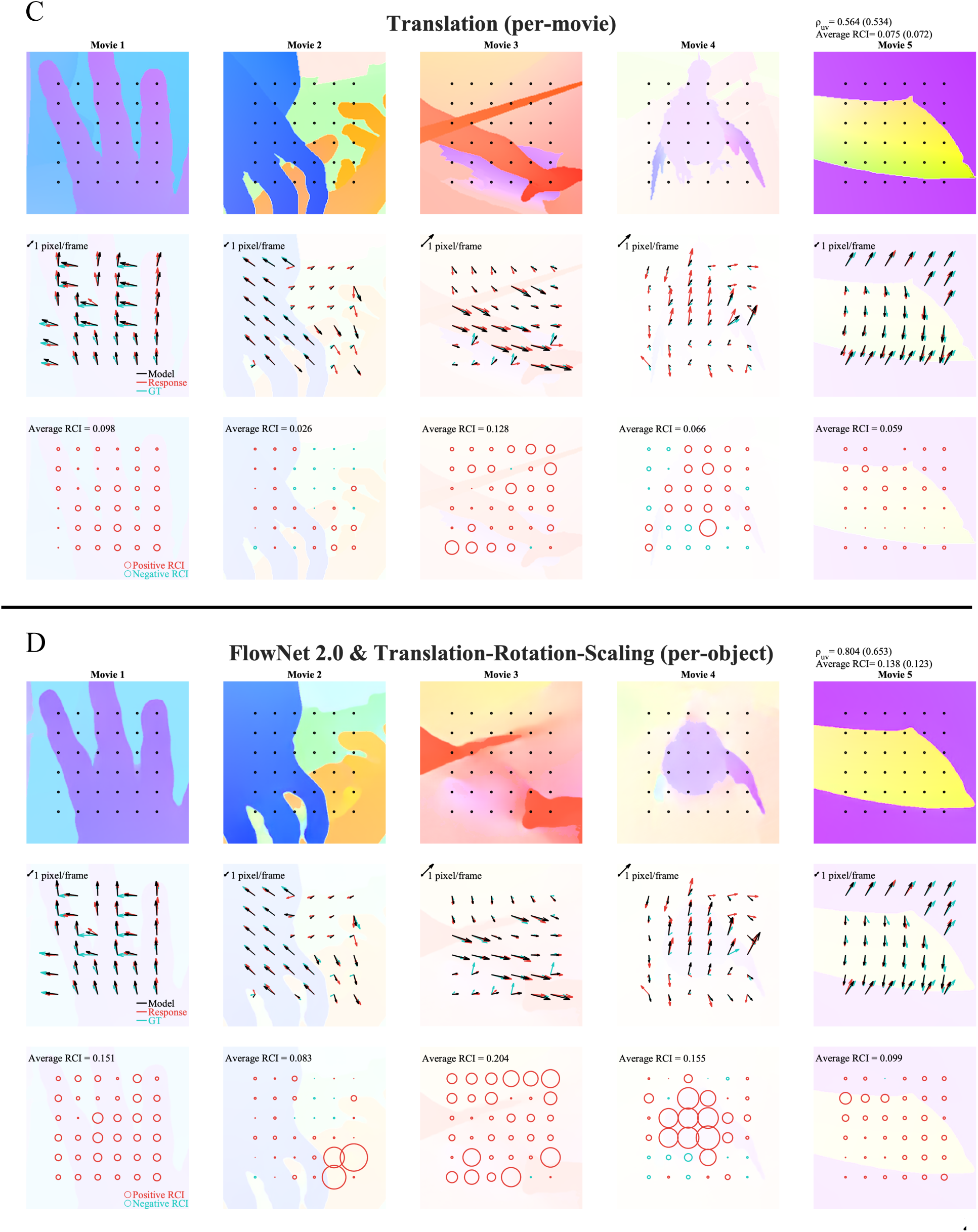
Spatial maps showing the extent of consistency between human responses and model predictions for (A) a pooling model (spatial pooling), (B) a computer vision model (FlowNet2.0), (C) a coordinate transformation model (Translation (per-movie)), and (D) a hybrid model (FlowNet 2.0 & Translation-Rotation-Scaling (per-object)) (see Fig. S9 for the results of all models tested). In each panel, the first row is the color-coded visualization of the optical flows predicted by each model. The second row depicts the motion vectors in each location; the black arrow denotes the model-predicted response, the red arrow the human response, and the green arrow the GT. The plotting scale (the length of a 1 pixel/frame vector) is shown by an arrow in the top left corner of each panel. The diameters of the circles in the third row indicate the RCIs, with red circles indicating positive values (the deviations from the GTs were consistent between the model predictions and the human responses) and green circles indicating negative values (the deviations from the GTs were inconsistent). The average RCIs of the 36 locations in each movie are shown in the top left corners.

### Low-level pooling models

Previous studies have shown that the spatial resolution of human visual motion processing is low. Detection of spatial modulations in optical flow is most sensitive at spatial frequencies lower than 1 c/°, becoming more difficult at higher frequencies (19, 20). The low spatial resolution may partially explain why perceived vectors deviate from the GTs. Although many neural mechanisms may contribute to the low spatial resolution of visual motion perception (21,12, 22), we modeled this simply via 2D Gaussian pooling of the GT vectors and estimated the amplitude and sigma of the Gaussian weighting function that best explained the human response. The standard deviation of the best-fit function was ∼5 pixels (Table S2). Small but positive cross-validated partial correlations with the human response (*ρ_uv_* = 0.239) and a positive averaged RCI (0.15) indicate that the spatial pooling model explains some aspects of the response errors. In agreement with our expectation that spatial pooling would be associated with deviations from the GTs at points close to motion boundaries, the points with high RCIs indeed lie near object borders (Fig. 5A). In addition, when points distant and close to borders were separately analyzed (see Fig. S9 for the definition of border points), agreement between the model and the responses was evident for only the border points (Table S4).

The temporal resolution of human visual motion processing is also low (23, 24). To model this, we performed 1D Gaussian pooling of the GT motion vectors over time and estimated the best amplitude, mean, and sigma of the Gaussian weighting function that explained the human response. The (cross-validated) partial correlations (*ρ_uv_* = 0.101) and averaged RCI (0.10) were low. When the spatial and temporal poolings were combined into a 3D Gaussian spatiotemporal pool, the power afforded in terms of explaining the human response was no better than that of the spatial pooling model (*ρ_uv_* = 0.214, average RCI = 0.15). The cross-validation-based statistical tests revealed that *ρ_uv_* did not differ significantly among the three pooling models (Fig. S7B), and that the average RCI was lower for the temporal pooling model than the other two models (Fig. S7D). Also, the high correlations among the spatial RCI patterns (Table S3) indicate that the three pooling models might explain similar aspects of the human response errors. It should be additionally noted that the movie clips tested here include only temporally smooth optical flows; the results might be different if stimuli featuring large temporal changes had been used.

### Computer vision models for optical flow estimation

Next, we compared the human responses to the outputs of five computer vision models developed for optical flow estimation. Farnebäck (25) is a conventional model that estimates dense optical flow. FFV1MT (26) is a biologically motivated model that includes a feedforward, primary visual cortex (V1)-middle temporal area (MT) structure. FlowNet 2.0 (27), StaRFlow (28), and RAFT (29) are convolutional neural net models based on supervised learning; they employ different architectures when seeking to improve GT estimations. Specifically, FlowNet 2.0 features multiple subnetworks, StaRFlow a spatiotemporal recurrent architecture, and RAFT iterative refinement of optical flow (see Supplemental Materials and Methods for the details).

Table 1 indicates that the RAFT model output was the closest to the human response in terms of the EPE and the correlations, but this was because the model output was also the closest to the GT. When comparing the partial correlation and the average RCI, FlowNet 2.0 best explained the human response errors, but the explanatory powers of the other computer vision models were only slightly lower (with the exception of StaRFlow). Bootstrapping-based tests indicated that *ρ_uv_* did not differ significantly between FlowNet 2.0, RAFT, Farnebäck, and FFV1MT (Fig. S7A), but the average RCI was higher for FFV1MT than RAFT (Fig. S7C).

The RCI spatial map of FlowNet 2.0 (Fig. 5B) reveals at which probe locations the model and humans make similar errors. Most such locations lie where dark objects make movements that differ from those of adjacent regions (see the lower right corner of Movie 2, and the lower part of Movie 3). It is difficult to estimate correct optical flows at such points because of a lack of image information (signal loss). The RCI spatial maps are similar for the various computer vision models, particularly between Farnebäck and FFV1MT and between FlowNet 2.0 and RAFT (Fig. S8, Table S3).

As compared to the pooling models, cross-validation-based tests on *ρ_uv_* and the average RCI indicated that the explanatory power for the human response was significantly higher for FFV1MT, FlowNet 2.0, and RAFT (Fig. S7 B, D). Also, the RCI spatial maps of the computer vision models correlated positively with those of the pooling models (Table S3), presumably because the former models include certain pooling processes.

### Coordinate transformation models

The pooling and computer vision models explain only some of the flow illusions (*ρ_uv_* ≤ 0.386). What remains to be explained? The models considered above process optical flow using only retinal image coordinates, as does early visual motion processing in humans. Certain illusions in terms of human visual motion perception, including induced motion (30, 31) and biological motion (13), may be associated with higher-level vector analysis that transforms optical flows from image coordinates to world- or object-centered coordinates. Allied phenomena include mutual repulsion/attraction between adjacent objects, such as motion contrast (32), direction repulsion (33), and motion capture (34). To evaluate the contributions of these higher-level optical flow illusions to human motion perception when viewing naturalistic scenes, we fitted the following four transformation models to our data.

In the Translation (per-movie) model, a common translation vector is subtracted from the GTs at all points in the same movie clip. This is one of the simplest descriptive models of induced motion and vector analysis (13); a motion pattern of multiple objects is decomposed into a global common motion of a reference frame and local relative motions. In the Translation-Rotation-Scaling (per movie) model, which tests more complex coordinate transformations, a common rotation (in direction) and rescaling (of speed), in addition to a common translation, are applied to the GTs of all points in the same movie clip. In the Translation (per-object) model, a common translation vector is subtracted from the GTs at all points in the same object layer of each movie. During such analysis, 36 points in each movie are classified into 2-3 groups depending on the object layer to which each point belongs (see Fig. S9 for how to define object layers). Separate parameter-fitting of different objects can capture flow distortions among multiple objects, including attraction and repulsion; these cannot be captured by global transformations. In the Translation-Rotation-Scaling (per-object) model, a common translation, a rotation in direction, and a rescaling of speed are applied to the GTs at all points in the same object layer of each movie. In all cases, the transformation parameters for each movie or object layer that best fit the human responses were estimated (see Table S2).

The four models captured the pattern of human errors rather well (*ρ_uv_* = 0.534-0.597, average RCI = 0.072-0.121). Compared to the low-level pooling and computer vision models, the coordinate transformation models showed significantly higher partial correlations (Fig. S7B) and average RCIs (Fig. S7D) when assessing human responses. Also, the RCI spatial maps differed from those of the low-level models. For example, the coordinate transformation models well-predicted the pattern of human errors in Movie 4 (Fig. 5C and S8C), probably because induced motion altered the perceived motion of a flying chicken.

When the four translation models were compared, cross-validation-based statistical tests on the average RCIs indicated that model complexity increased the explanatory power of the response data; the Translation-Rotation-Scaling models outperformed the Translation models, and per-object models outperformed per-movie models. However, statistical tests on *ρ_uv_* (Fig. S7B) indicated that only the difference between the Translation (per-movie) model and the Translation-Rotation-Scaling (per-movie) model attained statistical significance. A close inspection of the cross-validation results indicated that per-object models sometimes exhibited visible prediction errors at points close to object layer boundaries, possibly because of the lack of adequate numbers of points for the object layers and response ambiguities at object boundaries. These outliers affected the *ρ_uv_* more than the average RCI. From the data currently available, we conclude that subtracting a single global translation for each movie is too simple to fully describe the effect of perceptual transformation, but we require data that are more dense before we can conclude how much more complexity we must add. Note that the RCI spatial maps of the four models (Fig. S8C) are similar (see also Table S3), suggesting that the mechanisms underlying the perceived errors predicted by these models may be shared or overlap significantly.

The successful fitting of coordinate transformation models to human response data suggests that human perception of naturalistic movies is affected not only by lower-level effects but also by higher-level factors such as induced motion and vector decomposition.

### Hybrid models

If we are correct in stating that computer vision models reflect lower-order visual motion processing and transformation models higher-order motion processing, with each explaining different aspects of the human errors, it would be expected that combinations of such models would predict human errors better than either alone. We therefore computed the outputs of coordinate transformation models after changing the inputs from GTs to the optical flows estimated by FlowNet 2.0, which evidenced the highest *ρ_uv_* of all five tested models. As expected, we found that the *ρ_uv_* values and average RCIs for the hybrid models were better than those of the original transformation models employing GTs. The increases in both indices attained statistical significance for the two per-movie models (Fig. S7 B,D), but not for the Translation-Rotation-Scaling (per-object) model. In summary, as far as we tested on the current data, a hybrid model architecture best explains human-perceived optical flow.

## Discussion

Traditionally, vision scientists have sought to understand the mechanisms of human visual perception by analyzing relationships between subjective perceptual experiences and the responses of neural mechanisms and/or the predictions of computational models. Recent technical advances have enabled researchers to access large-scale data on neural responses and model behaviors for complex inputs including natural stimuli. Thus, data on human visual perception should become big and rich. Here, we psychophysically measured the human-perceived optical flows of dynamic naturalistic movie clips, using a novel method based on motion matching and flash probing. We first confirmed that our method was adequately accurate. However, the perceived flows of naturalistic stimuli deviated from the GTs in ways that depended on the stimulus structures (“flow illusions”). By comparing the human-perceived flows to the predictions of a variety of image-computable models, we concluded that the illusions were attributable to both lower-level factors such as spatiotemporal pooling and signal loss, and higher-level factors such as inter-object interactions and coordinate transformations resulting from vector analysis. Earlier studies showed that such factors produced specific illusions when certain experimental stimuli were presented to the observers, but we are the first to show that a combination of such factors explains the human-perceived optic flows of naturalistic scenes. Our method scales up the data on subjective visual experience, paving the way for a leap in human motion perception research.

To the best of our knowledge, human-perceived optical flow maps of naturalistic scenes have not been reported previously. There are several reasons. First, appropriate stimuli are lacking. When exploring how the visual system processes a stimulus of the external world embedded in images, one needs to know the GT of the variable. Many movies with natural scenes are available, but accurate GTs of the optical flows are not easy to derive. To overcome this problem, the computer vision community uses synthesized movies for training and evaluation of models, and the MPI Sintel Dataset is one of the most popular among such resources. We thus used this dataset.

The second difficulty is the workload. When estimating human-perceived flows, perceived motion vectors must be measured at many points. In psychophysical experiments, such measurements cannot be made in parallel, but must be made separately for each location. Use of a strict psychophysical procedure (e.g., estimation of the point of perceptual equality via a forced-choice paired comparison of direction or speed employing the method of Constant) would be very time-consuming. Thus, we derived a quick (thus practical) psychophysical method: motion vector matching by a method of adjustment. We limited the number of points to 180, sampled at 1° intervals within 5 x 5° regions of five movie clips. We measured the responses of four well-trained observers under four flipping conditions that effectively excluded response biases. Despite the “time-saving” measurements, the observers rather accurately reported vectors perceived at or around spatiotemporal positions indicated by the flash probe (SI 1). As various phenomena including flash lag (35) and flash drag (36) can cause (apparent) spatiotemporal misalignments between continuously moving and flashed patterns, we were concerned that our flashed probe method might not accurately identify spatiotemporal positions in moving patterns. It was therefore important to confirm that the spatiotemporal accuracy afforded by flash probing was high, at least for the temporally smooth motion clips that we used.

Another novel contribution of the present study is direct comparison of human-perceived flows with the flows predicted by computer vision models. Four of the five models that we tested, including FlowNet2.0, similarly explained some human response errors caused principally by input signal loss. Our model selection is not exclusive. Also, we did not fine-tune the parameters and/or structure of any model to match human performance. Therefore, we cannot conclude from the present data which computer vision model best simulates human visual motion processing. However, our study suggests that high-performance computer vision algorithms serve as proxies of an ideal observer model (37) that estimates retinotopic optic flows close to the GTs of input images with little internal noise.

The limited capacities of the computer vision models in terms of explaining human response errors may reflect differences in the computational goals. The models that we tested are aimed at estimating local image shifts accurately between frames in image coordinates. Human visual motion processing is aimed not only at estimating optical flows in retinal image coordinates but also at estimating object motions in appropriate coordinate frames, self-motions relative to the environments, and relative object depths as revealed by motion parallax. Based on optic flows estimated via lower-level processing, higher-level processing attain biologically meaningful goals when viewing complex natural stimuli, while produce motion illusions when viewing simple artificial stimuli. The fact that transformation models explain human responses well supports this view.

The ultimate goal of this project is to develop an image-computable model that can account for the perceived human flows for arbitrary visual inputs. One promising model architecture is a hybrid model in which optical flows estimated by image-based motion detectors are fed to higher-level processing, including coordinate transformation. The question then is how to predict the behaviors of the higher-level processing (e.g., coordinate transformation parameters) directly from the input image sequence. One strategy is the development of a computational model for high-level human motion processing (38). An alternative is a data-driven approach; artificial neural networks could be trained using human response data. In either case, it is necessary to scale up the data size, thus to collect more human responses at higher spatiotemporal densities with a broader range of stimuli.

We also expect our study will usefully connect psychophysics and neuroscience, since our data can be directly compared to the cortical activities of humans or animals measured via fMRI, optical imaging, or other methods, while observers watch the same movie clips. However, a few issues may arise when comparing our data to neural and behavioral data collected under different viewing conditions. We measured perceived motion vectors at single spatiotemporal points while directing each observer’s gaze and attention to those points. The perceived flow map of a collection of independent local measurements may significantly differ from a neural response map that is recorded in parallel at a time. Specifically, our procedure may underestimate context effects. Also, use of a random-dot motion field as a matching stimulus may direct the observer’s attention more to textural motion flow than to high-level object trajectories.

Finally, complex naturalistic stimuli and simple artificial stimuli are both useful for understanding visual processing (39). In our case, several visual illusions having been studied using artificial stimuli can explain a significant proportion of flow illusions arising during perception of naturalistic movies. This is quite reasonable based on the idea that many of the visual illusions arising when viewing simple artificial images should reflect visual processing that attempts to achieve higher-level visual functions (e.g., perceptual constancy) in complex natural scenes. Extending this idea, it could be argued that visualization and modeling of complex human perceptions of natural scenes, which we are challenging, is the ultimate way to understand the computational goals and mechanisms producing the conventional illusions triggered by simple artificial stimuli.

## Materials and Methods

This study was approved by the Research Ethics Committee of the Graduate School of Informatics, Kyoto University (approval no. KUIS-EAR-2020-003). Three of the authors and a naïve laboratory member participated in the experiments. All observers had normal or corrected-to-normal vision and gave informed consent before the study. Visual stimuli were presented on the display of a 13-inch MacBook Pro or a monitor controlled by a desktop Windows PC.

For the main experiment, we used five short movie clips selected from a slower version of the MPI Sintel Dataset (42) rendered at a frame rate of 1,008 frames per second (FPS) and a resolution of 2,048 x 872 pixels. We presented these clips at 60 FPS after magnifying the clip image to 4,098 x 1,744 pixels before clipping it to fit a 600-pixel circular aperture. In each clip, we defined 36 probed locations on a six-by-six grid with spacings of 50 pixels (1°) (see Fig. 1C and Movie S2). In each trial, the target motion clip was reframed such that the to-be-probed location was at the center of the aperture. Each movie clip was presented under four flip conditions (see Fig. 1E). Throughout the experiment, four dots (five pixels in diameter) were presented 60 pixels above, below, left, and right of the display center as position markers; these indicated to the observers the location of the display center to which they were to pay attention during the task. The five movie clips were tested in separate blocks, each consisting of 144 trials (36 flash-probed locations × four flip conditions). Each observer performed 720 trials.

We used a method of adjustment to measure the motion vector that the observer perceived at a specific spatiotemporal location in a short video clip (Fig. 1A). In each trial, a target motion stimulus and a matching stimulus were repeatedly presented on a uniform grey screen. The target clip was presented at the center of the display in a circular window (600 pixels or 12° in diameter). In the middle of the 15-frame target clip presentation (Fig. 1B), a probe dot (five pixels in diameter) was flashed at the display center for one frame (16.67 ms). The matching stimulus was a circular Brownian-noise field (120 pixels in diameter, 100% peak contrast) presented at the center of the display. The observers were asked to report the motion vector at the location/time indicated by the target probe by matching the speed and direction of the matching stimulus using the mouse cursor. When satisfied with the matching, the observer clicked the mouse, but early termination (before three repetition cycles) was not accepted. At the end of each trial, the speed and direction of the target and matching stimuli were shown as numbers on the display. This feedback was aimed at helping observers develop appropriate response strategies.

When comparing human responses and model predictions, we initially averaged the four human responses and the data for the four flip conditions in the original image coordinates, thus obtaining 180 data points (i.e., five movies x 36 positions) to contrast with vectors predicted by each model. To evaluate model performance, we computed average EPEs, Pearson correlations, partial correlations, and the RCI, which was the product of three terms, *A*·*B*·*C*, where:

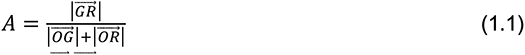

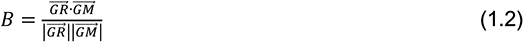

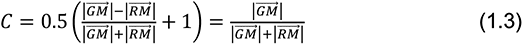

and G, R, and M are the endpoints of the GT, response, and model prediction vectors, respectively, and O the origin.

We used the least-squares fitting method to determine the best-fit parameters that minimized the summed EPEs of human optical flow (see Table S4 for all best-fit parameters). The numbers of free parameters were 2 (spatial pooling), 3 (temporal pooling), 4 (spatiotemporal pooling), 0 (computer vision models), 10 (Translation [per-movie], 2 x 5 movies), 20 (Translation-rotation-scaling [per-movie], 4 x 5 movies), 24 (Translation [per-object], 2 x 12 objects). and 48 (Translation-rotation-scaling [per-object], 4 x 12 objects). We used two-fold cross-validation (with common sampling sets) to evaluate statistically the explanatory powers of the models.

More detailed Materials and Methods information can be found in the Supplementary Information (SI3).

## Data availability

The data used (Dataset S1-S4) and stimulus movies (Movie S1-S2) can be accessed on https://doi.org/10.17605/OSF.IO/BU7PD.

## Acknowledgments

This research was supported by the Japan Society for the Promotion of Science (KAKENHI grants JP20H00603 and JP20H05957).

## Supporting Information Text

### SI 1: The accuracy of the proposed method in terms of estimating perceived optical flow

We evaluated two aspects of the accuracy of the proposed method. First, we explored how accurately the observers reproduced the direction and speed of the perceived flow (vector-matching accuracy). We analyzed the data of a preliminary experiment using a random-dot kinematogram and the data of the main experiment employing MPI Sintel movie clips.

We also evaluated how accurately (in space and time) the observers reported local vectors specified by the flashed probe (flash-probing accuracy). We analyzed the data of the main experiment. Specifically, we presented the movie clips under four flipped conditions (no flip, horizontal flip, vertical flip, and horizontal and vertical flip, see Fig. 1E) to decompose the two types of response error. One is an error defined by the display coordinates, thus determined by the relationship between the observer and the display, not by the image pattern. For example, an observer may tend to overestimate rightward motion or over-report vectors located left of the flashed probe. One can visualize errors of this type by analyzing the response vectors of the flipped conditions per se (i.e., without unflipping). We analyzed the responses on display coordinates to evaluate the accuracy of our method. The other type is response errors defined on the image coordinates whose directions flip as the images flip. One can visualize the errors of this type by unflipping the four flip conditions to the original image coordinates.

#### Vector-matching accuracy of the Random-Dot Kinematogram (RDK)

In a preliminary experiment, we measured the human responses to RDKs with spatiotemporally uniform optical fields. We used the same method as that used for the MPI SINTEL experiment; we changed only the target stimulus. The RDK stimulus consisted of 5,000 white and black dots at a density ratio of 50-50 on a gray background. Each dot was 3 pixels (3.6 arcmins) in diameter, and the total dot density was ∼20%. Note that this traditional random-dot pattern differs from that of the Brownian matching-noise stimulus. All dots moved coherently in one direction at a constant speed. The speed and direction were randomly and uniformly selected from 1–10 pixels per frame (PPF) and 0–360° respectively. The four position-marker dots and the probe dot were red. There were two blocks, each with 144 trials. Each observer underwent 288 trials. To evaluate accuracy in terms of reporting the probed vector, we examined the relationship between the reports and the GTs in terms of direction, and the speed and horizontal and vertical (*uv*) components of the vector (Fig. S1). In terms of direction, the responses of the four observers (288 trials) were very accurate. The coefficient of determination for the linear regression (square of the Pearson correlation coefficient) was *r^2^* = 0.998, and the coefficient of determination for the ideal response model *R^2^* = 0.998. *R^2^* refers to the proportion of variance in the human response (*Resp_i_*) that can be predicted by the GT:

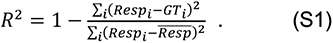

In terms of speed, the response was more scattered and the regression trended to the mean (*r^2^* = 0.641, *R^2^* = 0.533). The observers reported speeds consistent with those of the presented stimuli, although the accuracy was not as high as that for direction. The *uv* vector components, which combine direction and speed information, evidenced high positive correlations between the GTs and the responses (*r^2^* = 0.957, *R^2^* = 0.901). In summary, the results of the preliminary test suggested that the procedure enabled observers to report reliably the speed and direction of perceived motion.

#### Vector-matching accuracy for the MPI Sintel Flow Dataset

As we did for the RDK data, we analyzed the accuracies of the direction and speed reports for the perceived flows of the naturalistic motion stimulus set. The scatter plots of the responses versus GTs using the raw data (i.e., five movies × 36 locations × four flip presentations ×four participants) for the direction, speed, and *uv* components are shown in Figure S2A-C. The vectors are shown in the display coordinates (i.e., the vectors of the flipped conditions are not unflipped to the original orientations). The response accuracy measures were *r^2^* = 0.860 and *R^2^* = 0.859 for direction, *r^2^* = 0.466 and *R^2^* = 0.333 for speed, and *r^2^* = 0.657 and *R^2^* = 0.643 for the *uv* components. The agreements between the reported and GT vectors were reduced in the main MPI SINTEL experiment compared to the preliminary RDK experiment.

Our principal interest involved the question of why the vector-matching accuracy for the naturalistic movie clips was reduced. Particularly, we sought constant response errors, or biases, associated with the stimulus image structure, not with the general traits of observers. To explore how stimulus-dependent factors affected response accuracy, we averaged the human responses over the four flipped conditions after unflipping the vectors into the original image coordinates for all four participants. The scatter plots of the averaged data (i.e., five movies × 36 locations) by the GT angles in the original image coordinates are shown in Figure S2D-F. The averaged response accuracy measures were *r^2^* = 0.932 and *R^2^* = 0.931 for direction, *r^2^* = 0.810 and *R^2^* = 0.648 for speed, and *r^2^* = 0.854 and *R^2^* = 0.836 for the *uv* vectors.

#### Flash-probing accuracy for the MPI Sintel Flow Dataset

If observers accurately reported vectors at the probed locations on the display, the response vectors would be expected to be closer to the GT vector at that location than at any other image location. Figure S3B shows a 2D map of the endpoint L2 distance between the response vector for a probe at (x, y, t) = (0, 0, 0) and the GT vector at (x, y, 0) where t = 0 refers to the central (eighth) frame. As in Figure S2A, the vectors of the four flipped conditions are represented in the display coordinates (i.e., without unflipping to the original orientation). Figure S3A shows a similar 2D map where the response vector is replaced with the GT vector at the probed position. The curve in Figure S3C shows slices of the two maps across lines passing the points of the minimum endpoint difference between the response and probed points shown by the dotted lines in Figure S3A and S3B. The point of minimum difference (1.031 at [-3, 0, 0]) is very close in terms of both position and amplitude to the probed point (1.034 at [0, 0, 0]). Note that the spatial unit is the pixel, and 1 pixel corresponds to 1.2 min arc. Although the response endpoint difference (red curve) fell gradually around the probed point, the similarity in the shapes of this curve and the GT endpoint difference curve (black) indicates that within-stimulus correlation, rather than inaccurate spatial localization by the observers, is the principal cause of the gradual drop. Human response variability explains the minor smoothing of the central peak of the response endpoint difference curve compared to the GT curve. As we merged the data for the four flip conditions in the display coordinates, the black curve is symmetrical. The fact that the red curve is also nearly symmetrical implies that the display coordinates exhibit little response bias (see Fig. S4 for a similar analysis of the image coordinates). The results thus suggest that the observers reported the probed vector fairly accurately in space (Fig. S3C).

Next, we considered temporal probing accuracy. Figure S3D shows how the response endpoint difference (red curve) and the GT (black curve) changed over time. The response (or GT endpoint difference) is the L2 distance between the response or the GT at (x, y, t) = (0, 0, 0) and the GTs at (0, 0, t). The response is minimal at the probed frame and runs near-parallel to the black curve; this reveals the within-stimulus correlations associated with temporal flow smoothness. This indicates that the observers reported probed vectors accurately over time.

### SI 2: Response consistency across observers

Figure S5 shows the response flow map for each observer, and Table S1 the Pearson correlations among observers, and the partial correlations among observers with the effects of GT removed. The Pearson correlation between observers ranges from 0.933 to 0.959 for direction, from 0.620 to 0.842 for speed, and from 0.805 to 0.891 for the *uv* components, indicating that the responses were consistent across the observers. In addition, the partial correlation with the effects of GT excluded among observers ranges from 0.301 to 0.590 for direction, 0.024 to 0.512 for speed, and 0.115 to 0.526 for the *uv* motion vectors, suggesting that the GT error patterns were positively correlated across the observers.

### SI 3: Supporting Methods and Materials

#### Ethics Statement

This study was approved by the Research Ethics Committee of the Graduate School of Informatics, Kyoto University (approval no. KUIS-EAR-2020-003). The experiments were performed in accordance with the Declaration of Helsinki, with the exception of preregistration.

#### Participants

Three of the authors (YY, TF, and ZS) and a naïve laboratory member (ZZ) participated in the experiments. One author (ZS) collected his data before he joined the research team. All observers had normal or corrected-to-normal vision and gave informed consent before the study.

#### Apparatus

All experimental codes were written in Python 3 (40) using PsychoPy (41). Visual stimuli were presented on the display of a 13-inch MacBook Pro (resolution 1,440 x 900, refresh rate 60 Hz, gamma 2.2, background gray luminance 150 cd/m^2^, viewing distance 57cm) in a normally illuminated room, or on a monitor (Eizo CG303W, 1,920 x 1,200, 60 Hz, 2.2, 45 cd/m^2^, 72 cm) controlled by a desktop Windows PC in a darkened chamber. For each setup, the viewing distance was adjusted so that 50 pixels subtended a visual angle of 1° to the observer.

#### Stimuli

In the main experiment, we did not use the standard MPI Sintel dataset (24 FPS, 1,024 x 436 pixels) but, rather, a higher-frame-rate version (42) originally rendered at 1,008 FPS and a resolution of 2,048 x 872 pixels. We selected five short movie clips that contained relatively complicated spatial structures and multiple objects. We magnified the clips to 4,098 x 1,744 pixels before clipping them to fit a 600-pixel circular aperture and presented them at 60 FPS. This enabled spatial sampling of reasonably dense perceived flow in a local image area while suppressing long-range jumps between adjacent frames, for which human motion detection may rely more on high-level feature tracking than low-level motion detection (SI ref. 1-3). In each clip, we defined 36 probed locations on a six-by-six grid with a spacing of 50 pixels (1°) (see Fig. 1C). The grid location was chosen so that the 36 probed points covered a variety of figure objects and the background. In each trial, the target motion clip was relocated such that the to-be-probed location came to the center of the aperture (Movie S1). Throughout the experiment, four dots (each five pixels in diameter) were presented 60 pixels above, below, left, and right of the display center (as position markers) to indicate the location of the display center to which attention was to be paid during the task. To render the flashed probe dot and the surrounding four position marker dots equally salient, regardless of the movie context, we computed the HSV values of the dot locations in the movie clip and imparted the reverse HSV values to the dots. Each movie clip was presented under four conditions: no flip (original orientation), horizontal flip, vertical flip, and horizontal and vertical flips (Fig. 1E). This allowed us to decompose response errors defined in the display and image coordinates (SI 1). Most visual motion illusions (e.g., induced motions) are determined by the relative relationships of image components, and we thus analyzed responses in the image coordinates by averaging the response vectors of the four flipped conditions after unflipping them to the original coordinates (e.g., Figs. 3, 5). The five movie clips were tested in separate blocks, each consisting of 144 trials (36 flash-probed locations × four flip conditions). The blocks were ordered randomly, and each observer underwent 720 trials in total.

#### Procedure

We used a method of adjustment to measure the motion vector that the observer perceived at a specific spatiotemporal location in a short video clip (Fig. 1A). During each trial, a target motion stimulus and a matching stimulus were repeatedly presented on a uniform grey screen. One cycle of repetition consisted of the target clip (250 ms, 15 frames), an interstimulus interval (ISI, 750 ms), the matching stimulus (250 ms), and an ISI (750 ms). The target clip was presented at the center of the display in a circular aperture (600 pixels or 12° in diameter). In the middle of the target clip presentation (i.e., in frame 8 of 15 frames), a probe dot five pixels in diameter was flashed at the display center for 1 frame (16.67 ms). The matching stimulus was a circular Brownian (1/f^2^) noise field (120 pixels in diameter, 100% peak contrast) presented at the center of the display. The broadband Brownian noise aided observers to perceive a wide range of image speeds without aliasing attributable to undersampling in time. To reduce the abrupt stimulus onset/offset of the target and matching stimuli, the stimulus contrast was linearly increased during the first half of the presentation, attained full contrast in the eighth frame, and decreased linearly in the latter half of the presentation. Observers were asked to report the motion vector at the location/time indicated by the target probe by matching the speed and direction of the stimulus using the mouse cursor. This was a purple dot (five pixels in diameter) freely moveable within a central circular area (600 pixels in diameter). Depending on the distance of the mouse cursor from the display center (from 0 to 300 pixels in radius), the speed of the matching stimulus varied logarithmically from 0 to 20 pixels per frame (PPF). When satisfied with the matching, the observers terminated the trial by clicking the mouse, but early termination before completion of three repetition cycles was not accepted.

At the end of each trial, the speed and direction of the target and the matching stimuli of the trial were shown as numbers on the display. One difficulty associated with the method of adjustment was the setting of an appropriate criterion for trial termination. It was likely that too-early termination would sacrifice accuracy, but that longer observation would not necessarily improve performance. The main purpose of feedback was to help observers develop appropriate response strategies.

#### Evaluation of model predictions

To compare the human responses with model predictions, we initially averaged all four human responses and the four flip conditions in the original image coordinates, yielding 180 data points (i.e., five movies x 36 positions) to contrast with the vectors predicted by each model (except when we evaluated accuracies using the display coordinates). To evaluate performance, we computed average EPEs, Pearson correlations, and partial correlations for each model (Table 1). The visual processing models included parameter-fitting, but the numbers of free parameters differed among the models. To ensure fair comparisons, the values estimated via two-fold cross-validation were also calculated. A bootstrapping method was used to simulate the population distributions of the indices. Additionally, we generated RCIs to evaluate how the models predicted human response errors at each location.

##### Endpoint Error (EPE)

The EPE is the Euclidean distance between the GT (*μ_GT_,ν_GT_*) and human response (*μ_Resp_,ν_Resp_*) motion vectors as shown in equation (S2):

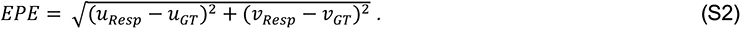

##### Comparison of direction

When we compared correlations in direction (r and ρ), we evaluated the shortest angular distance of the angle of the response, or the model (*θ_Resp,or model_*), from the angle of the GT (*θ_GT_*), by adjusting the phase of *θ_Resp,or models_* as follows:

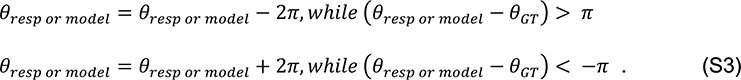

##### Partial Correlation *ρ*

For each model, we calculated partial correlation coefficients between the predicted outputs and the human response by controlling for the effect of GT to determine which fitting functions/computer vision models best explained the pattern of deviations in the human response from the GT.

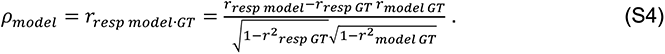

##### Response consistency index

The RCI is the product of three terms, *A*·*B*·*C*, where:

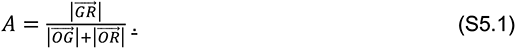

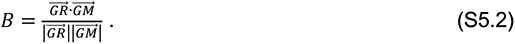

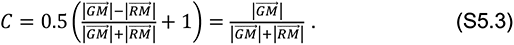

and G, R, and M, are the endpoints of the GT, response, and model prediction vectors respectively, and O is the origin. A highlights points of interest (i.e., those evidencing flow illusions). A is 0 when the GT and the response are in perfect agreement 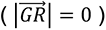 but 1 when they completely disagree 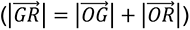. B specifies the direction similarity (the cosine of the direction difference angle) between the response error relative to the GT 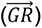 and the model error relative to the GT 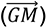. B is 1 when agreement is perfect, and -1 when there is complete disagreement. C compares the distance between the model prediction and the GT 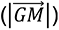 and the distance between the prediction and the response 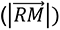. C is 1 when the prediction agrees with the response 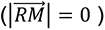 but 0 when the model agrees with the GT 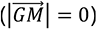. When the terms are combined, the RCI becomes positive and approaches 1 at locations where the model predicts conspicuous flow illusions; the RCI becomes negative at locations where the model predicts illusions in the opposite direction. This index is similar to the partial correlation between the model predictions and responses with the effects of GT removed (*r_dir_* = 0.886, *r_spd_* = 0.909 and *r_uv_* = 0.974 when non-cross-validated partial correlations are used), but the index evaluates model predictability at each location with integrating the direction and speed components.

##### Cross-validation

Two-fold cross-validation (SI ref. 4) was employed to evaluate model fitting statistically. During each cross-validation repetition, the 180 probed locations were split into two halves, thus subsets X and ∼X, in a constrained random manner (see below). The model predictions for X were computed based on the best-fit parameters for ∼X, and the model predictions for ∼X computed based on the best-fit parameters for X. Evaluation index values such as *ρ_uv_* were computed from the combined set of predictions for all 180 points. This was repeated 1,000 times to estimate the population distribution of each index value. Such analysis enables the direct comparison of models with different numbers of free parameters. For models with no free parameters, the estimated index values were always the same (we thus used a bootstrapping method to evaluate computer vision models statistically).

For the statistical comparison of the index values between models, the same cross-validation was used to compute the population distribution of differences in each index value between each pair of models. To exclude the effects of random subset selection on variation in the index value difference, the same sets of X and ∼X were used for all models during every cross-validation repetition. If the 95% confidence interval of the index value difference did not include zero, we regarded the difference between the models as statistically significant (alfa = 0.05, two-sided).

During cross-validation of the per-movie and per-object transformation models, we added a constraint to the random selection of subset X such that the locations were split into two halves within each movie and within each object layer (or approximately so when the number of points was odd). Except when otherwise noted, we used such constrained subset selection for all models and the same sets of X and ∼X for comparisons across the models, as noted above. We found that this constraint exerted only very minor effects on the estimated index values.

##### Bootstrapping

A bootstrapping method (SI ref. 5) was employed to estimate the population distributions of performance index values for each model, and, during statistical comparison of the models, the distributions of the differences in index values between each model pair. The fitting models employed the best-fit parameters for the response data at all probed locations (without cross-validation). During each of 1,000 repetitions, we re-sampled 180 locations from the original 180 locations with replacement, and computed the index values of each model and the differences among models. As during cross-validation, when comparing the models, we used the same re-sampling sets. Re-sampling was random with the constraint that the number of re-sampled data points was the same as the original number of locations within each object layer. We found that this constraint exerted very minor effects on the estimated index values.

#### Parameter-fitting Models

We used the least-squares fitting method to determine the best-fit parameters, minimizing the sums of human optical flow EPEs (see Table S4 for all best-fit parameters).

##### Spatial Pooling

This fitting function spatially aggregates local GT flow vector fields (*μ*_GT_,*ν*_GT_) to obtain a predicted perceived vector (*μ*_est_,*ν*_est_). A 2D Gaussian kernel *g*_S_(*x*,*y*) parameterized by the amplitude (*A*) and sigma (*σ*_S_) is used to weight the GT vectors surrounding the probed location (*x*_P_,*y*_P_). We did not consider the offset between the center of the Gaussian kernel and the probed location because the EPEs were symmetrically distributed between the human response and the flow vector field in 2D space (Fig. S3C). Therefore, the vector components (*μ*_est_,*ν*_est_) for each (*x*_P_,*y*_P_) were:

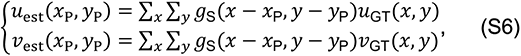

where:

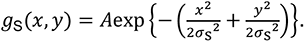

To reduce computational complexity, we constrained the spatial extent of the 2D Gaussian kernel to -30 to +30 pixels on both the x and y axes, with the probe at 0.

When separately analyzing the effects of spatial pooling on border and non-border areas, we categorized 55 of the 180 probed locations as border points, as indicated by the bracketed numbers in Figure S9, depending on whether a window of ±2.5*σ*_S_ around that point included more than one object layer or not, where *σ*_S_ was the best-fit sigma (5.025 pixels).

##### Temporal Pooling

This fitting function temporally aggregates GT flow vectors at the probed location. A 1D Gaussian kernel *g*_T_(*t*) parameterized by the amplitude (*A*), mean (*μ*_T_), and sigma (*σ*_T_) is used to weight the GT vectors from *t* = -7 to +7 frames (*t* = 0 refers to the probed frame) as:

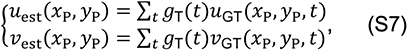

where:

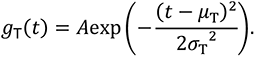

##### Spatiotemporal Pooling

This fitting function spatiotemporally aggregates the GT flow vectors. A 3D Gaussian kernel *g*_ST_(*t*) parameterized by the amplitude (*A*), spatial sigma (*σ*_S_) temporal mean (*μ*_T_), and temporal sigma (*σ*_T_) is used to weight the GT vectors as

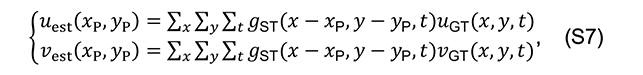

where:

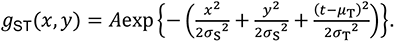

We constrained the spatial extent of the Gaussian kernel to -30 to +30 pixels on both the x and y axes, and the temporal extent to -7 to +7 frames.

##### Coordinate Transformations

The translation transformation decomposes the GT vectors, including the global translation vector (*μ*_0_,*ν*_0_), to parameters that fit the human response. The predicted perceived vector for each position is computed as follows:

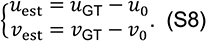

The translation-rotation-scaling transformation includes additional scaling (*s*) and rotation (*α*) parameters that affine transforms GT vectors to fit the human response:

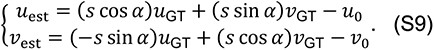

##### Per-Object Transformation Models

The per-object transformation models served as proxies of optical flow object-based processing. We manually defined object layers for each movie. In these models, we classified the 36 probed locations in each movie into layers with two or three objects and backgrounds based on their boundaries and the unique motion flows. The object layers are numbered as shown in Figure S9. Movie 1 contained a hand and background layers. Movie 2 contained left and right hands, and background layers (three layers). Movie 3 contained arm, animal, and background layers. Movie 4 contained a rooster and background layers. Movie 5 contained a bat’s wing and background layers.

#### Computer vision models

##### Farnebäck

This model uses a polynomial expansion transform to estimate the speed and direction of dense optical flows by approximating neighboring pixels between two frames based on the classic constant brightness assumption (I(x+uδt,y+vδt,t+δt) = I(x,y,t)) and global flow smoothness constraints (25). We estimated the displacements at five levels of the image pyramid via three iterations per level, using a 15×15 average window size and a Gaussian weighting function of SD = 1.1 for averaging over the neighborhoods. Each pixel neighborhood used to find polynomial expansions contained five pixels.

##### FFV1MT

This model is biologically inspired, and includes V1 motion energy estimations employing spatio-temporal Gabor filters and normalization, static nonlinear pooling of feedforward V1 responses in the MT layer, and a velocity estimate derived by decoding a linear weighed average of the MT response (26). The model also includes non-linear filtering of the MT response to handle spatial flow discontinuities more effectively. We selected -2∼2 subframes from the probed frame using the parameters of the original study (ref. 26, Table 2) and the code on the group site (SI ref. 6).

##### FlowNet 2.0

This is a convolutional neural network-based motion model that includes iterative refinement via multiple applications of FlowNetSimple (to extract motion vectors directly from two stacks of paired images) and FlowNetCorr (to extract motion vectors from a correlation layer for comparison of two separate images), to handle large displacements. FlowNet-SD is used to fine-tune the model with a focus on small displacements, and a final fusion network estimates motion flows (27). We trained the FlowNet 2.0 model using the Flyingchair, ChairsSDHom, and 3DFlyingthings datasets.

##### StaRFlow

This is a temporally dynamic, convolutional neural network-based motion model that includes spatio-temporal recurrent cells to generate multi-frame optical flow estimations and handle occlusion information (28). We trained this model on the MPI Sintel dataset (without experimental stimuli). Four continuous frames covered each training iteration with a batch size of 8, and the model attained convergence after 5,0000 iterations.

##### RAFT

This deep network motion model includes a feature encoder to extract features from two frames, a context encoder to extract context features from the first frame, a 4D correlation layer that computes the visual similarities of all pairs of pixel features at the 1/8 level, and an updating Gated recurrent unit to refine optical flow iteratively (29). Following the original study, we used the official model pre-trained on Flying chair and KITTI, and then fine-tuned the MPI Sintel dataset (without experimental stimuli) over 100,000 iterations with a batch size of 8.

---

**Fig. S1.**
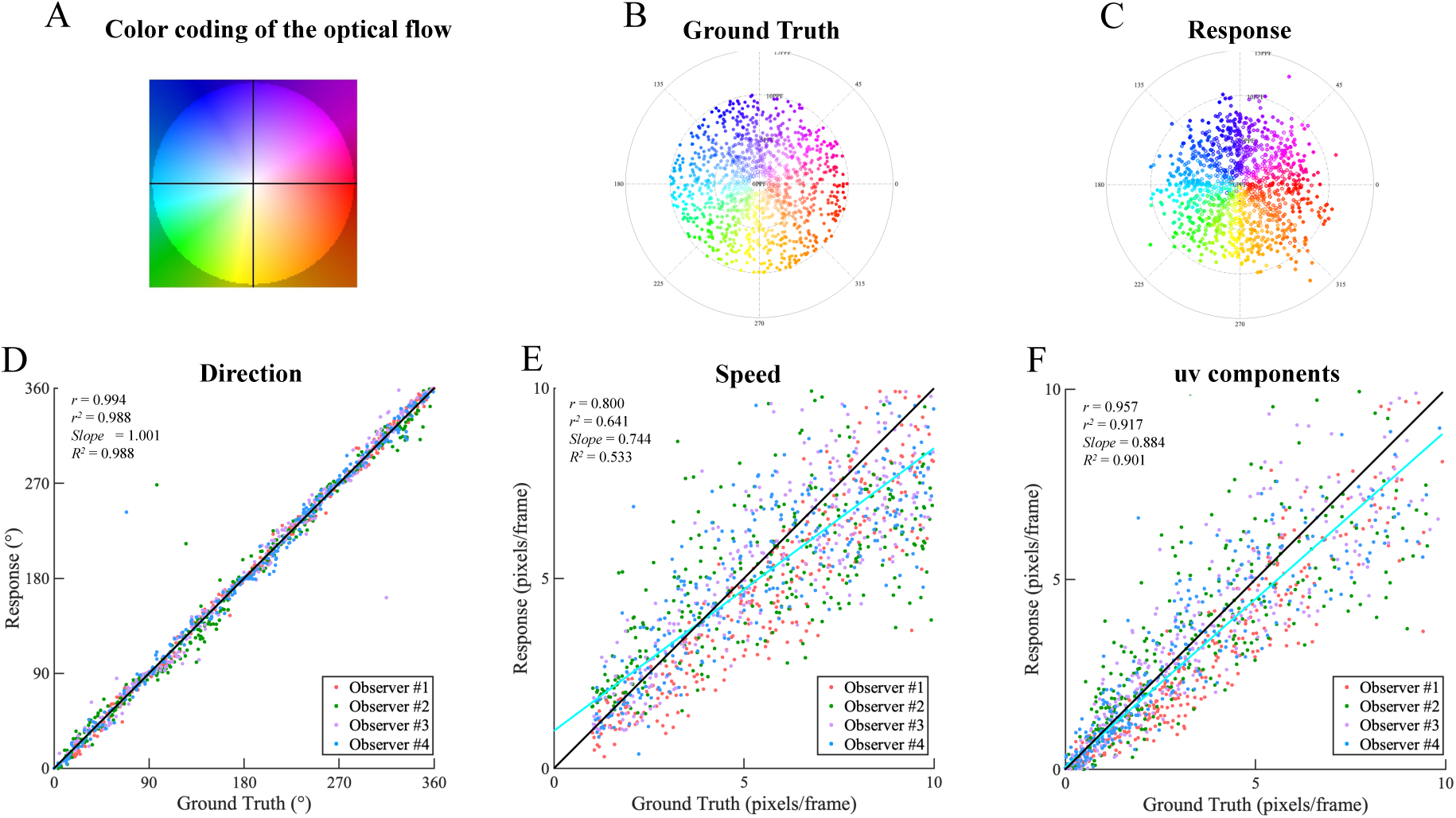
The results of the preliminary experiment using RDKs. (A) Color-coding is used to depict the GT optical flows of movie clips, with hue indicating motion direction and saturation motion magnitude. Polar plots of the vector distributions for (B) the GT and (C) the human response. The scatter plot of the response versus the GT for the moving direction (D), moving speed (E), and the *uv* components (F). Different colors refer to different individuals. *r*, *r^2^*: Pearson correlation and the square thereof; *slope*: the slope of the linear regression line; *R^2^*: the coefficient of determination by reference to the ideal response (Response = GT).

**Fig. S2.**
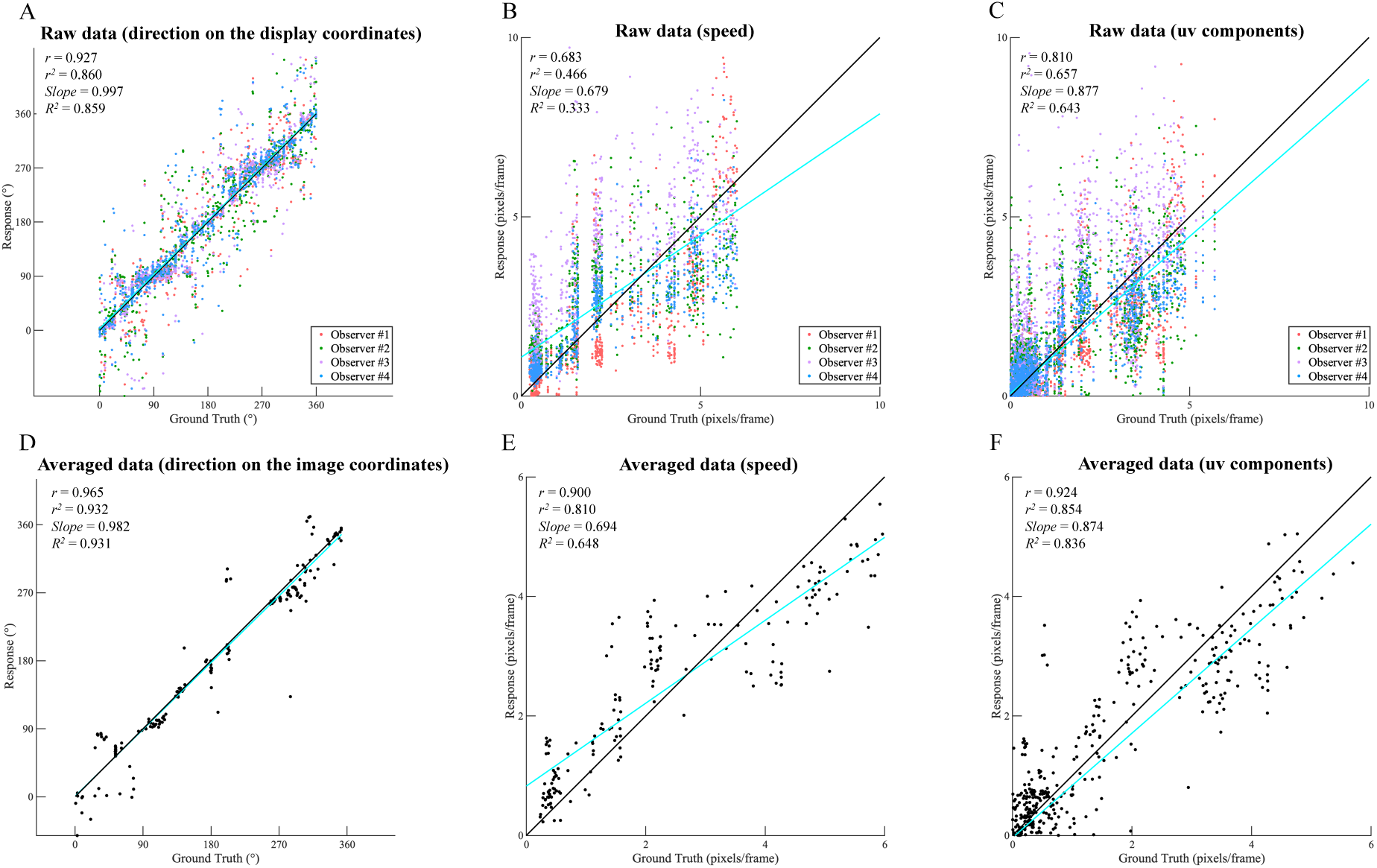
(A-C) The scatter plot of the human responses versus the GTs in the main MPI Sintel experiment. Raw data with the vector directions shown in the display coordinates for (A) the moving direction, (B) the moving speed, and (C) the *uv* components. Different colors refer to different individuals. (D-F) Data averaged over the individuals and the flip conditions with the vector directions in the original image coordinates for (D) the moving direction, (E) the moving speed, and (F) the *uv* components.

**Fig. S3.**
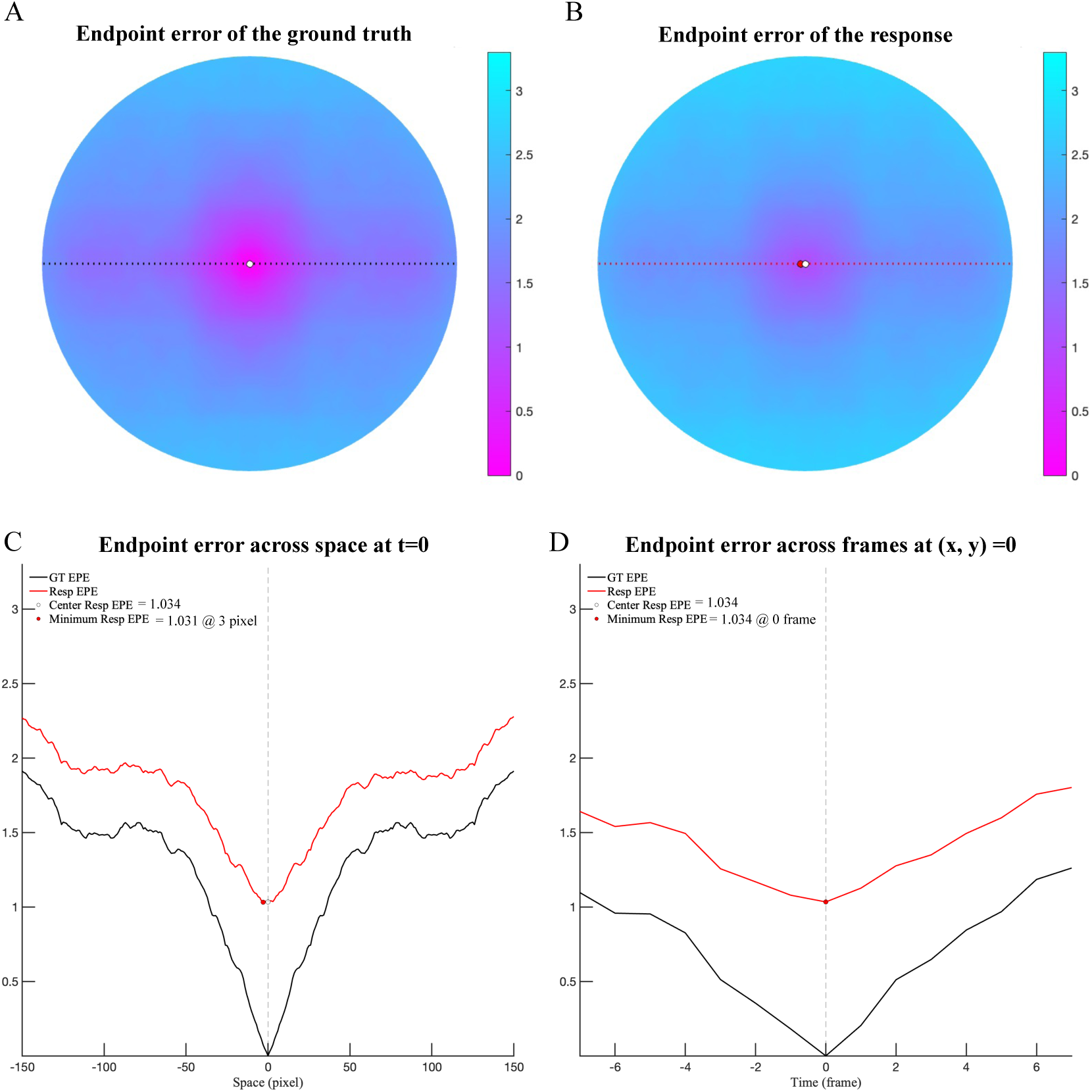
Flash-probing accuracy as shown by the spatiotemporal distributions of EPEs using the display coordinates of the MPI Sintel Flow Dataset. A 2D map of the endpoint L2 differences between (A) the GT at the probed location (0, 0, 0) and the GTs at all locations in the flashed frame (x, y, 0) (left) or (B) between the human response to a probe at (x, y, t) = (0, 0, 0) and the GTs of all locations in the flashed frame (x, y, 0). (C) The sectional profile (A, B) along the dashed line connecting the minimum difference location and the probed point. Note that the GT errors are spatially symmetrical. (C) The temporal profile of the endpoint differences. The red lines indicate the differences between the human response to the probe at (0, 0, 0) and the GTs at (0, 0, t). The minimum endpoint difference lies in the probed frame. The black lines indicate the endpoint difference between the GT at (0, 0, 0) and the GTs at (0, 0, t).

**Fig. S4.**
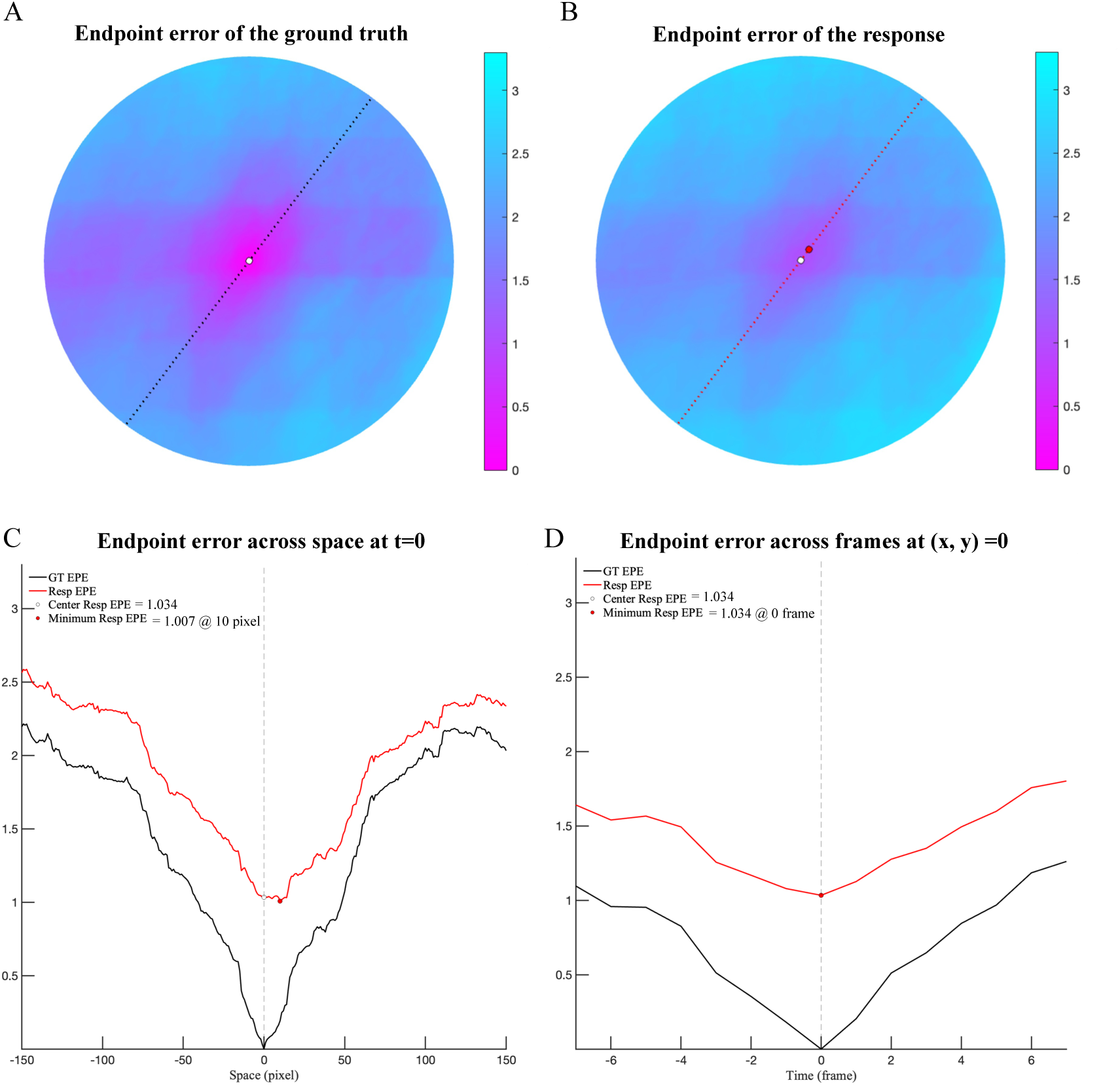
This is similar to Figure S3 except that the EPEs were computed using the original image coordinates. Note that the GT errors are not spatially symmetrical.

**Fig. S5.**
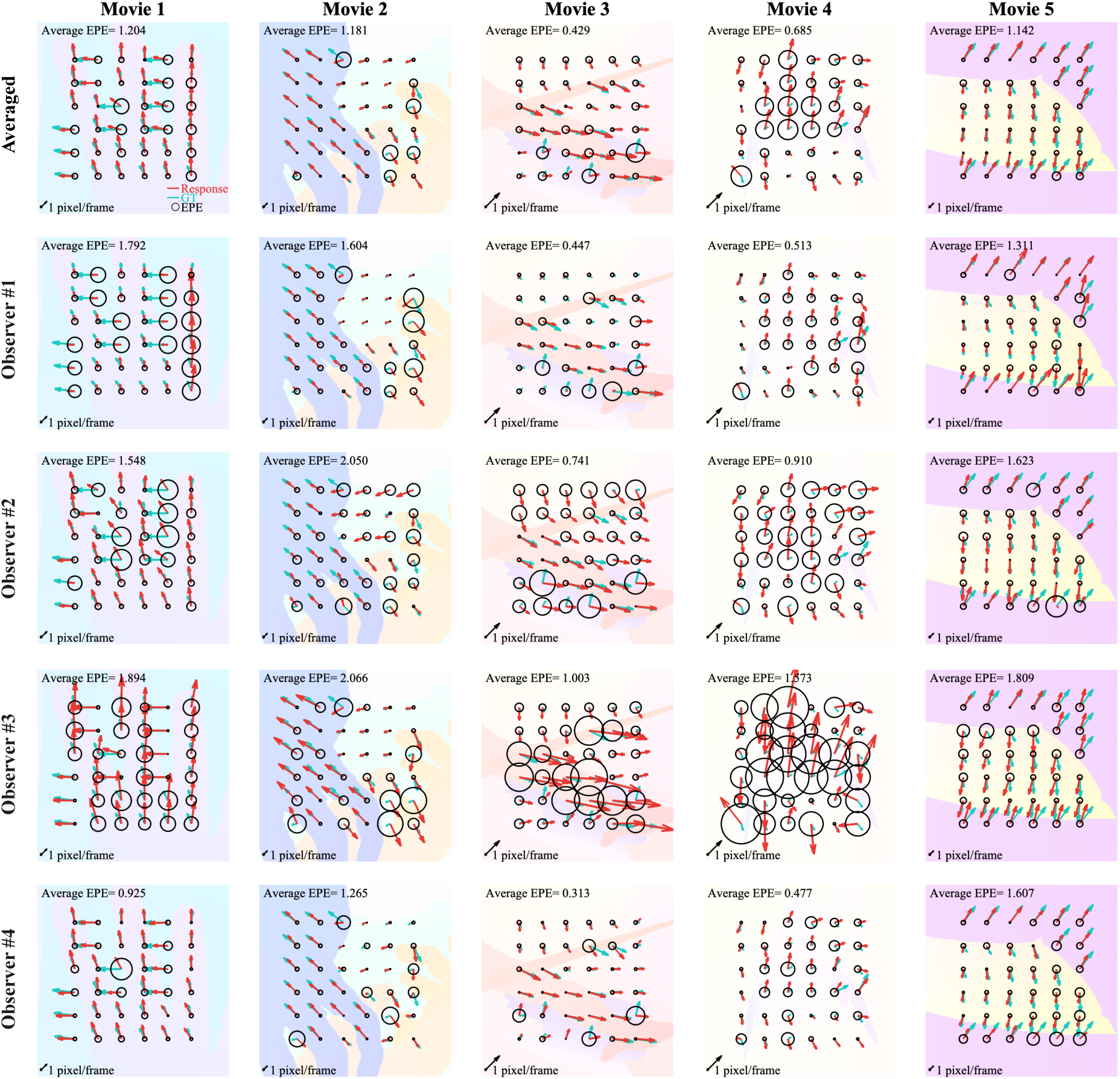
The response flow map in image coordinates for the MPI Sintel Flow Dataset. At each location, the green arrow denotes the GT, the red arrow the human response, and the diameter of the black circle the EPE. In each panel, the average EPE of the movie is shown in the top left corner. As the vector length is normalized within each movie, the spatial scale is shown by an arrow (of length 1 pixel/frame) in each bottom left corner. The first row indicates the response averaged over four observers, and the second to fifth rows the data of the individual observers.

**Fig. S6.**
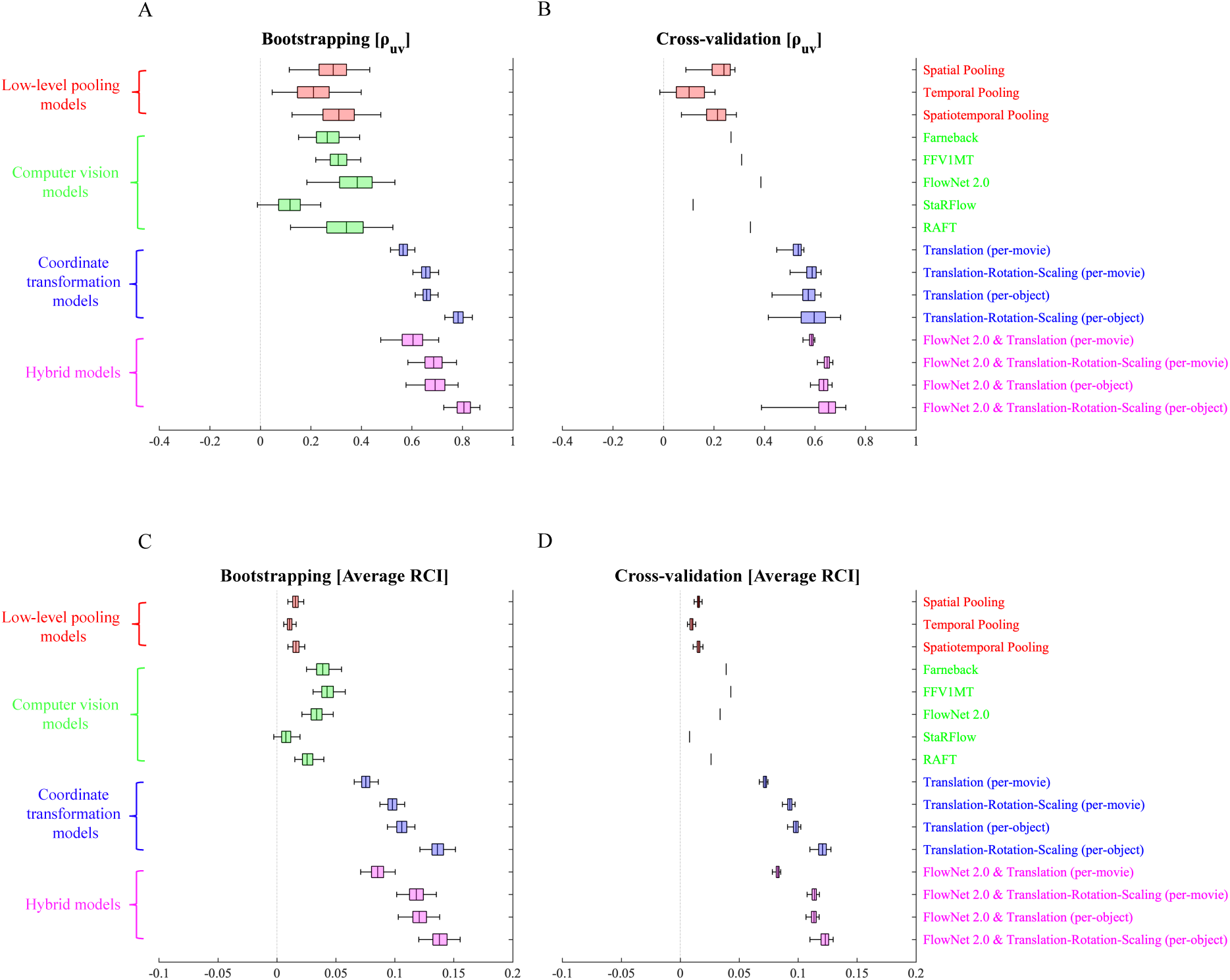
Box plots of the partial correlations *uv* components (ρ_uv_) and the average RCIs with the [25% percentile, median, 75% percentile] ranges (boxes) and 95% confidence intervals (whiskers). The ranges were sampled via (A & C) bootstrapping and (B & D) two-fold cross-validation. Note that the computer vision models were not subjected to cross-validation; the figures are those of the simple best-fit models.

**Fig. S7.**
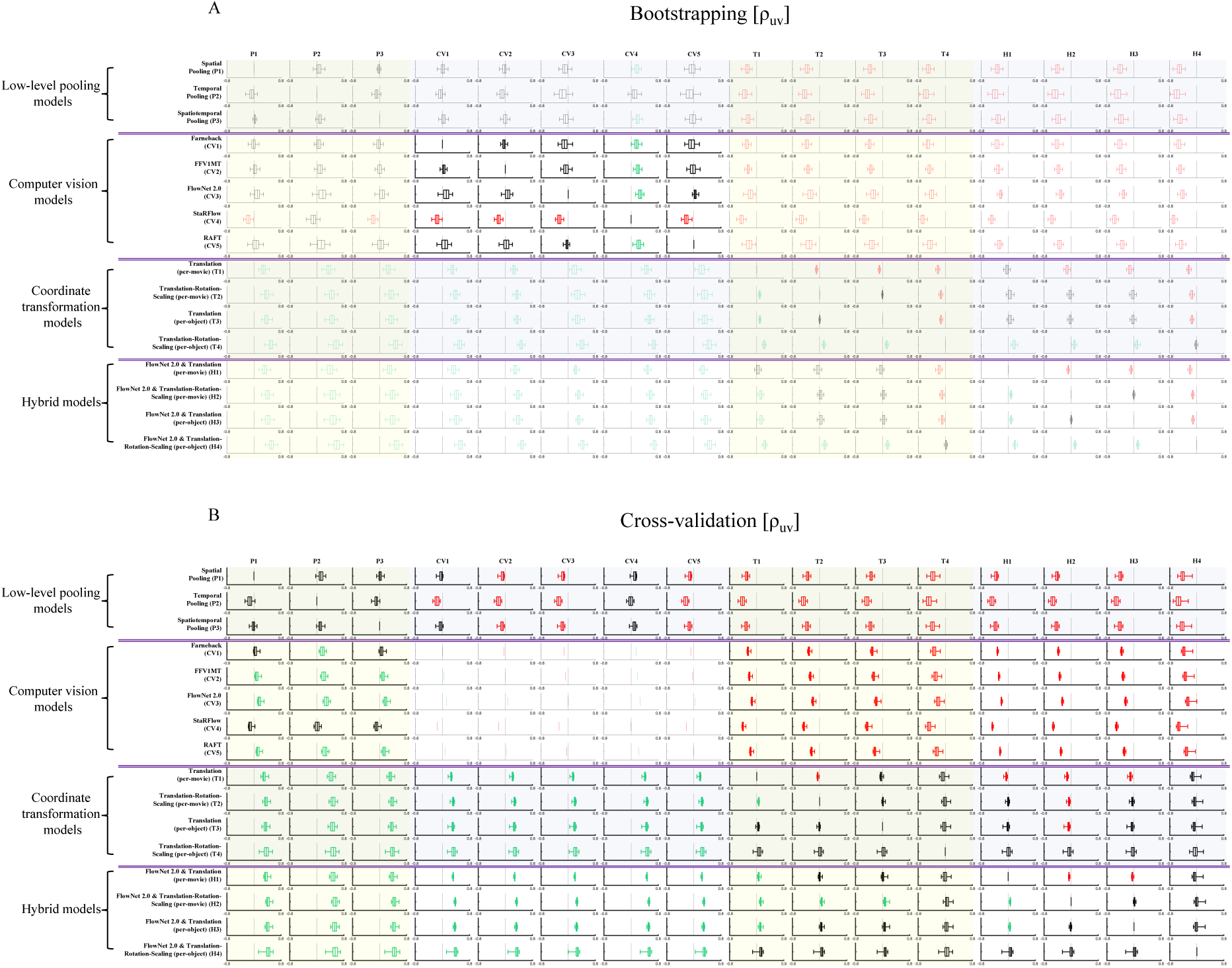
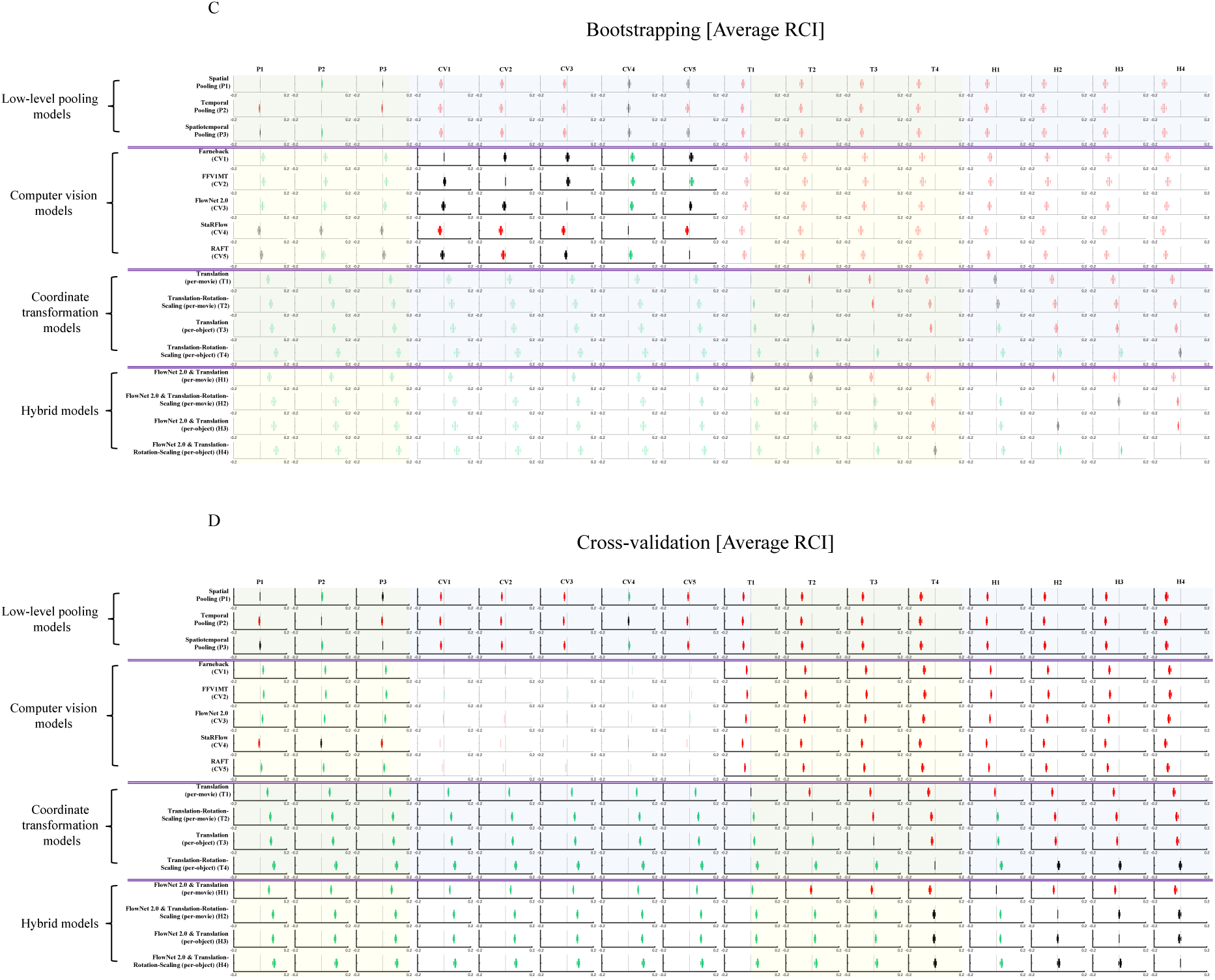
Pairwise model comparisons of the partial correlations *uv* components (ρ*_uv_*) and the average RCIs. The box plots show the differences between models in the horizontal row and those in the vertical column with [25% percentile, median, 75% percentile] ranges (boxes) and 95% confidence interval (whiskers). The ranges were estimated via (A & C) bootstrapping and (B & D) two-fold cross-validation. As computer vision models lack free parameters, the index value for the cross-validation was always the same as that of simple prediction. A green box indicates that the index value is significantly higher for the model in the row than the column in the sense that the confidence interval exceeds zero. A red box means the opposite; the index value is significantly lower for the model in the row.

**Fig. S8.**
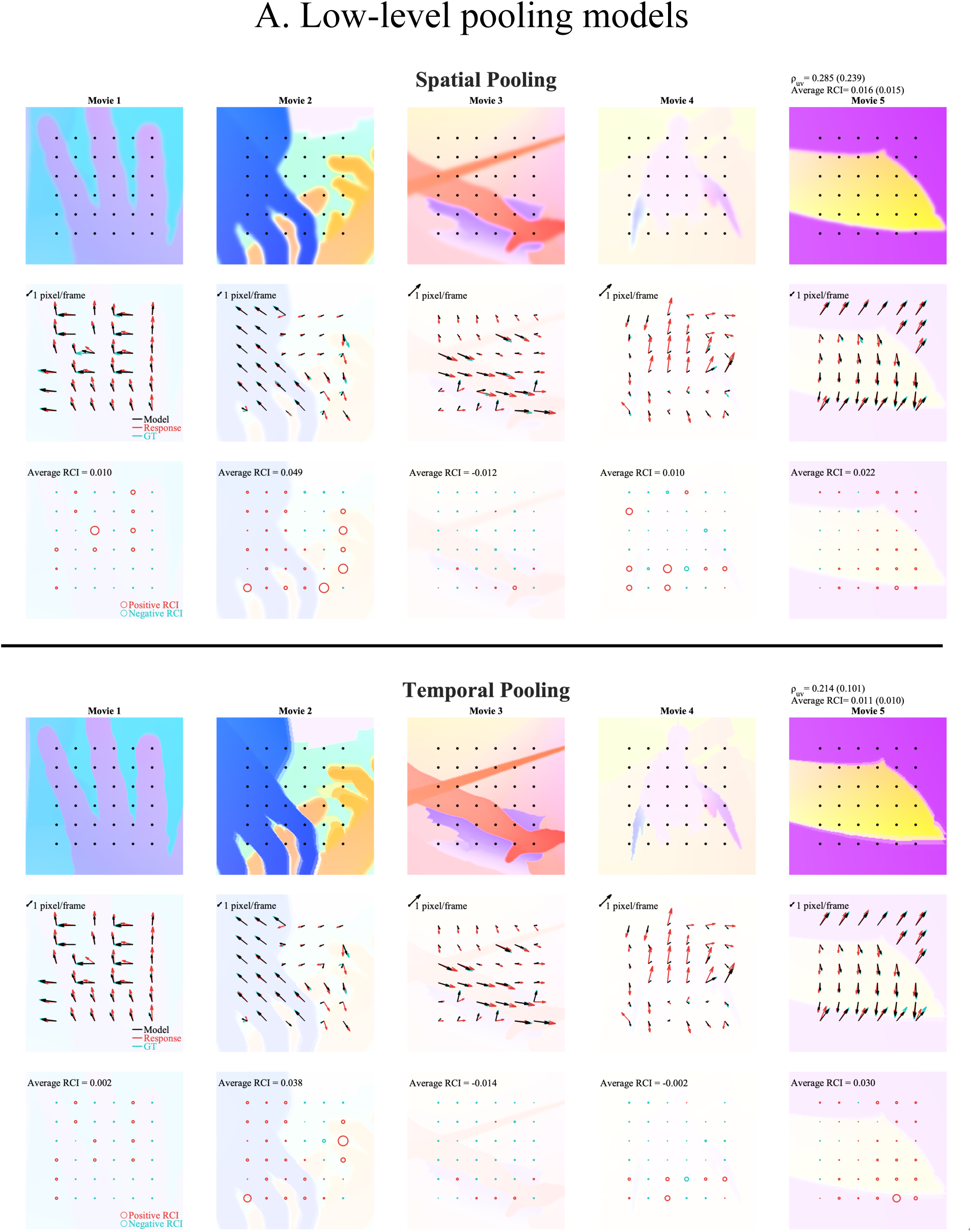
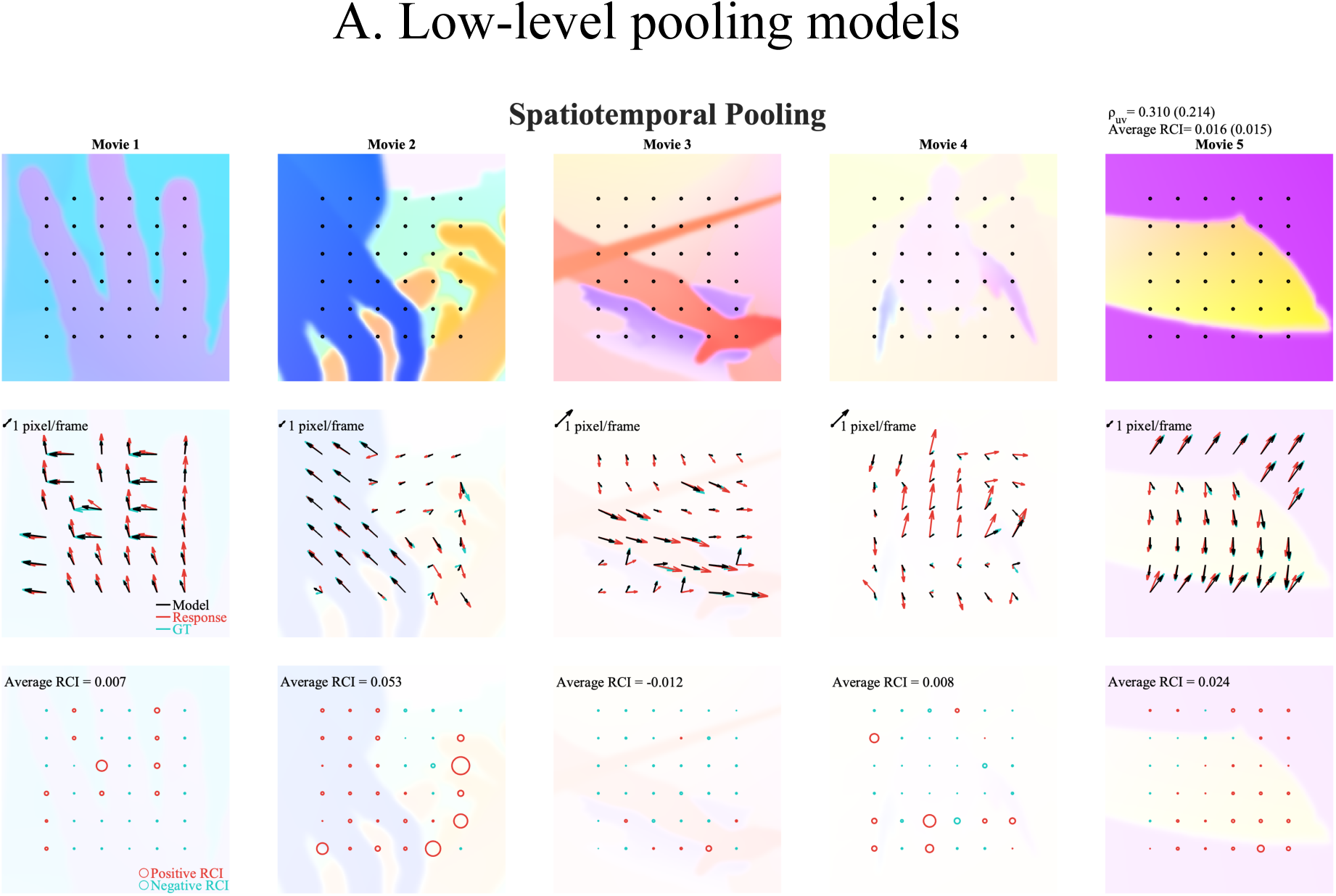
Spatial maps showing the consistencies between human responses and the model prediction for all models tested, in the same format as that of Figure 5. (A, this page) Low-level pooling models, (B) Computer vision models, (C) Coordinate transformation models, and (D) Hybrid models.

**Fig. S9.**
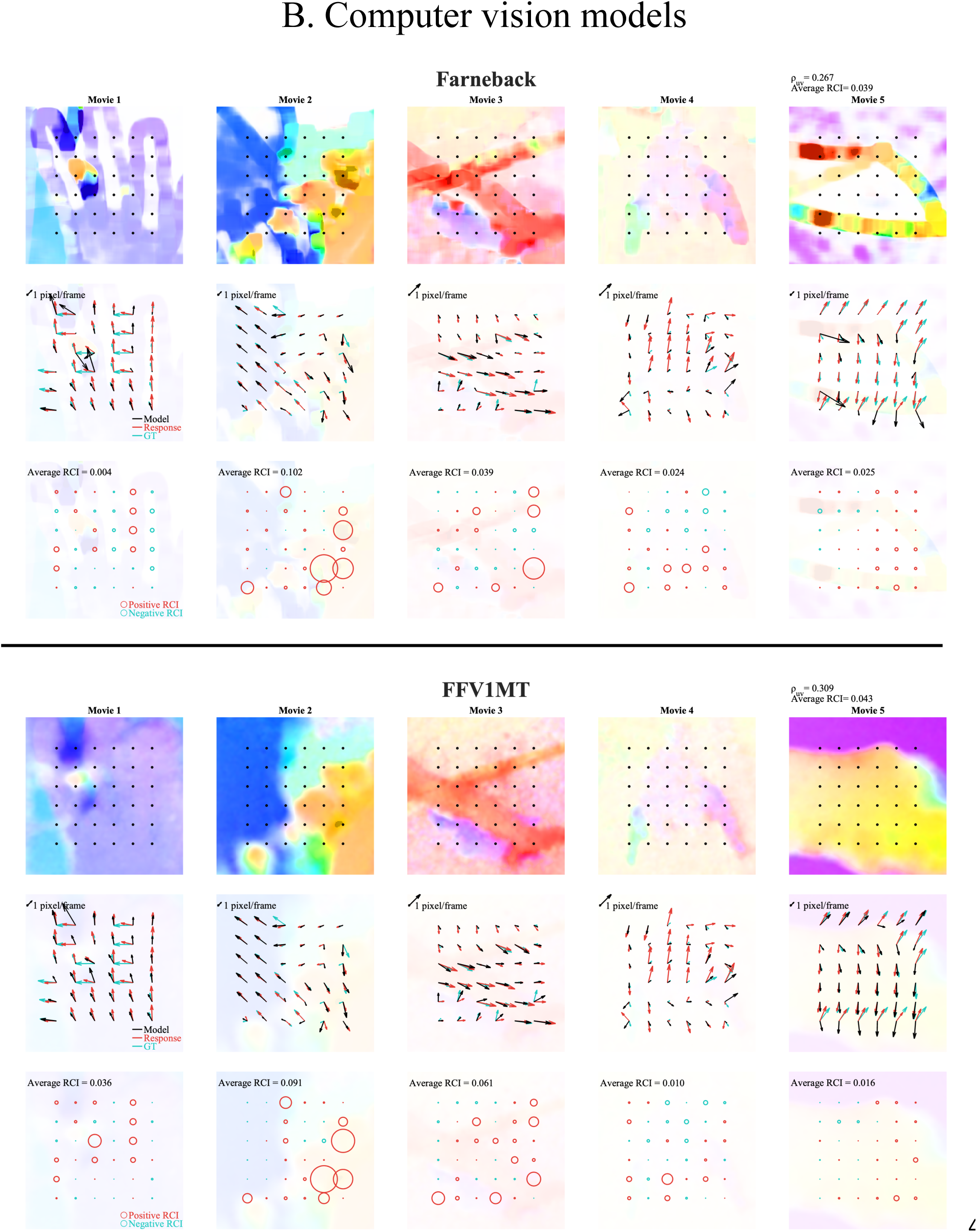
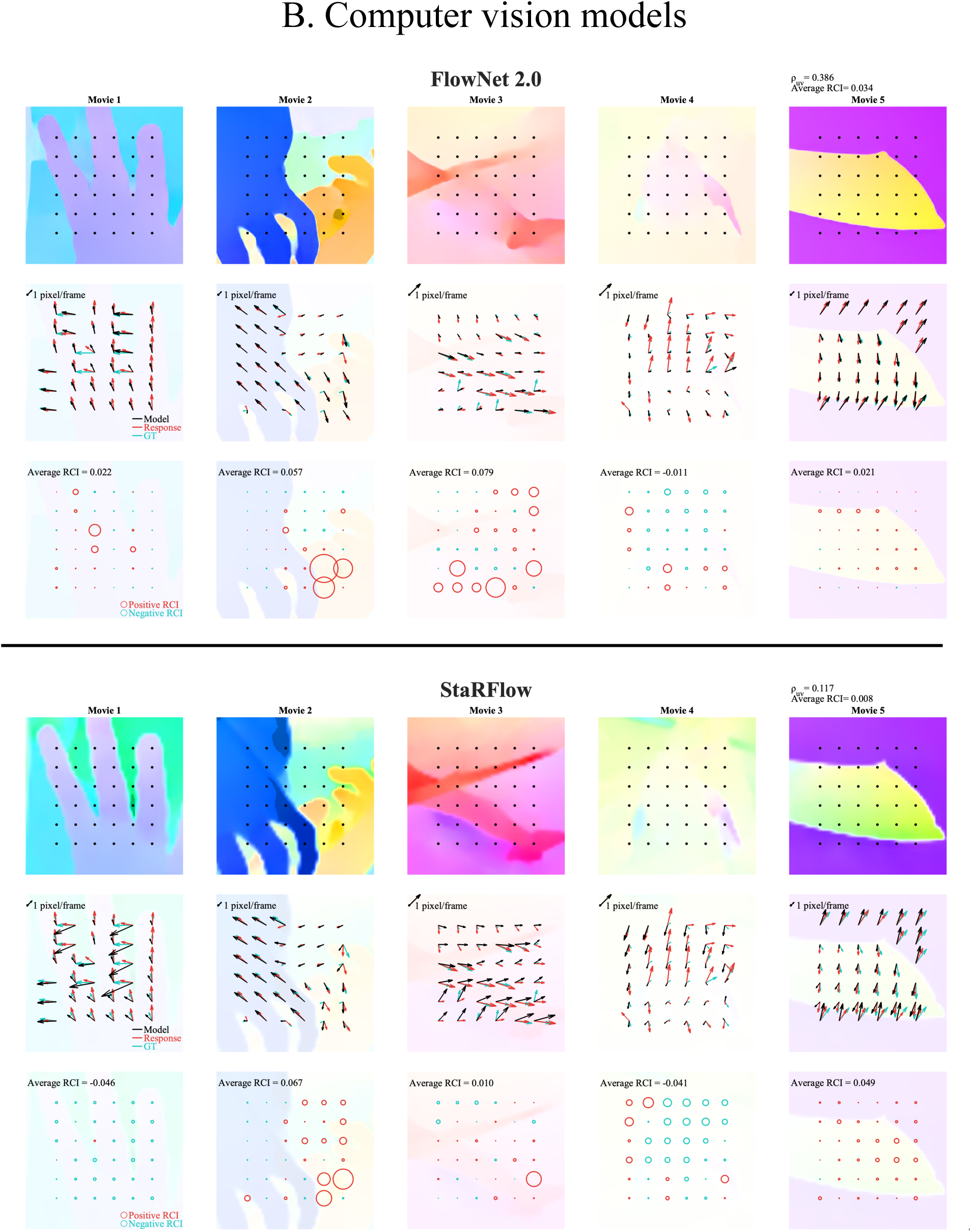
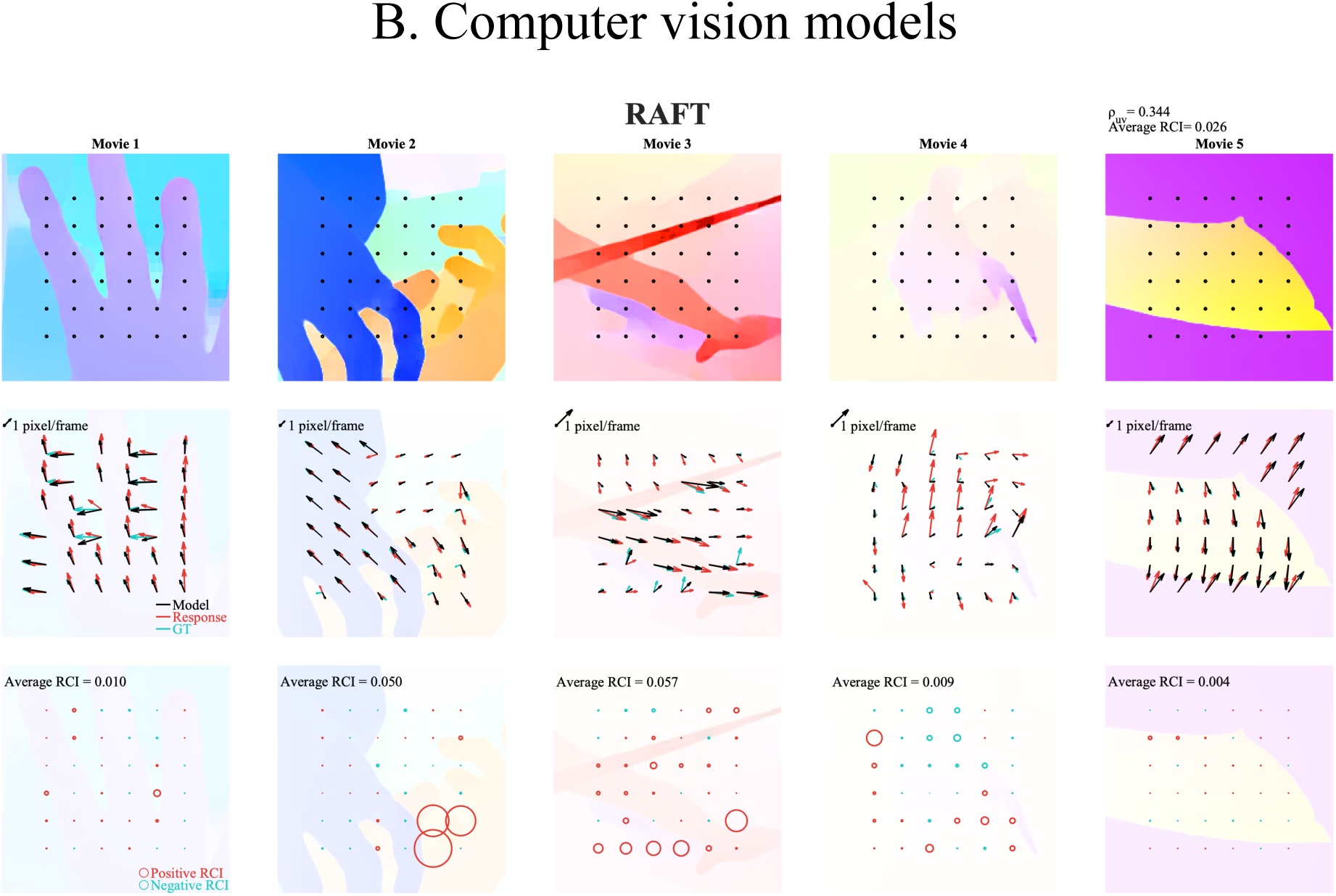
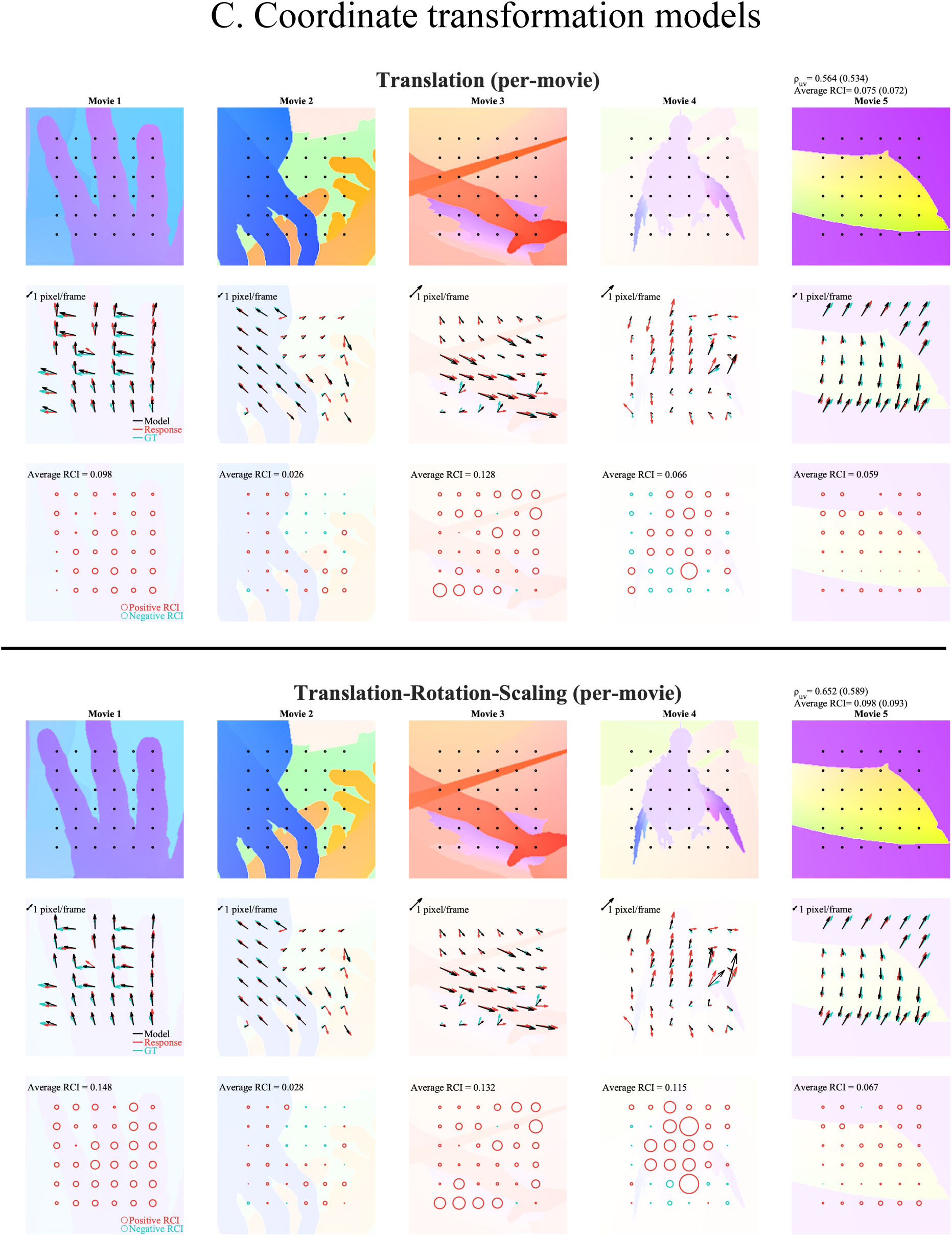
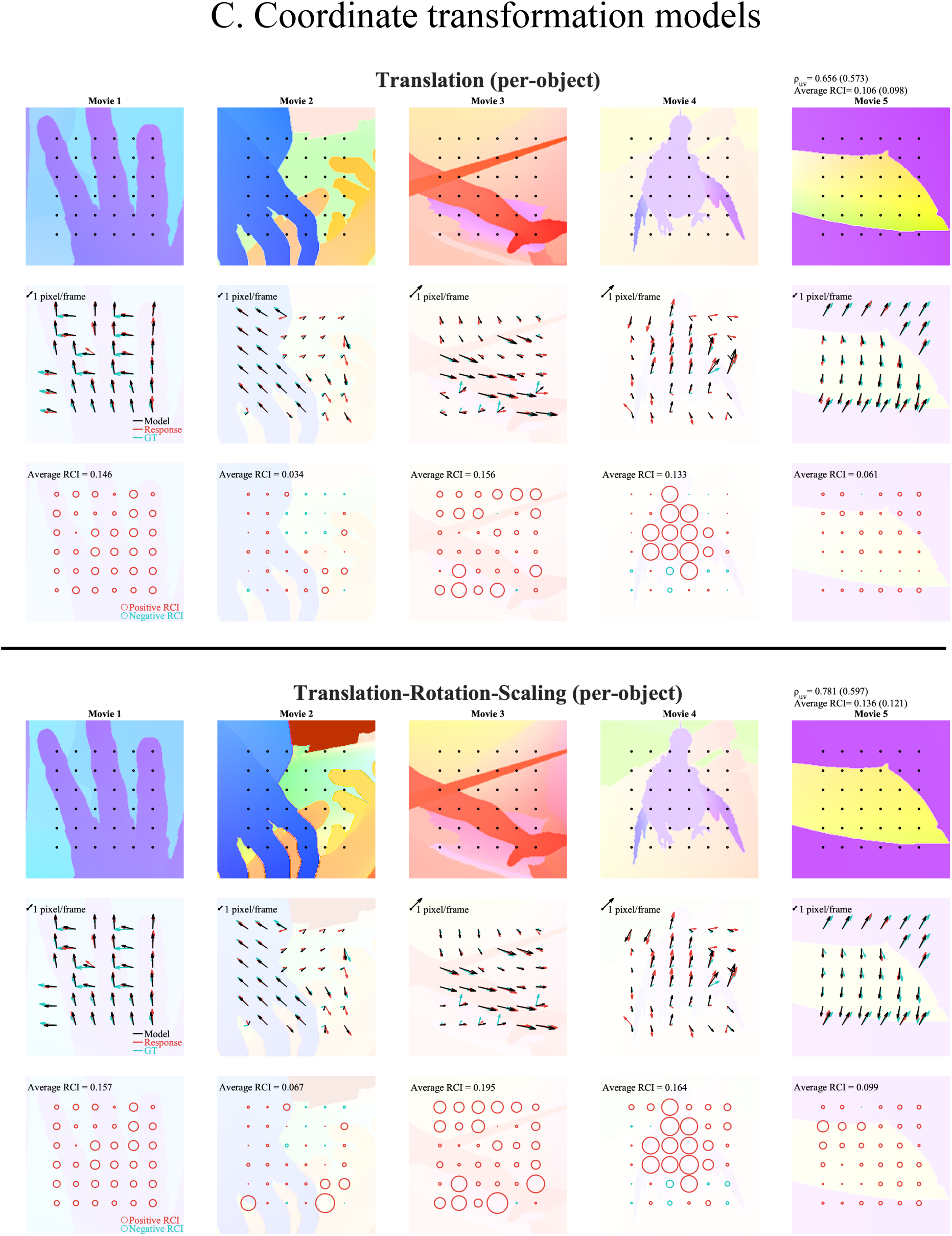
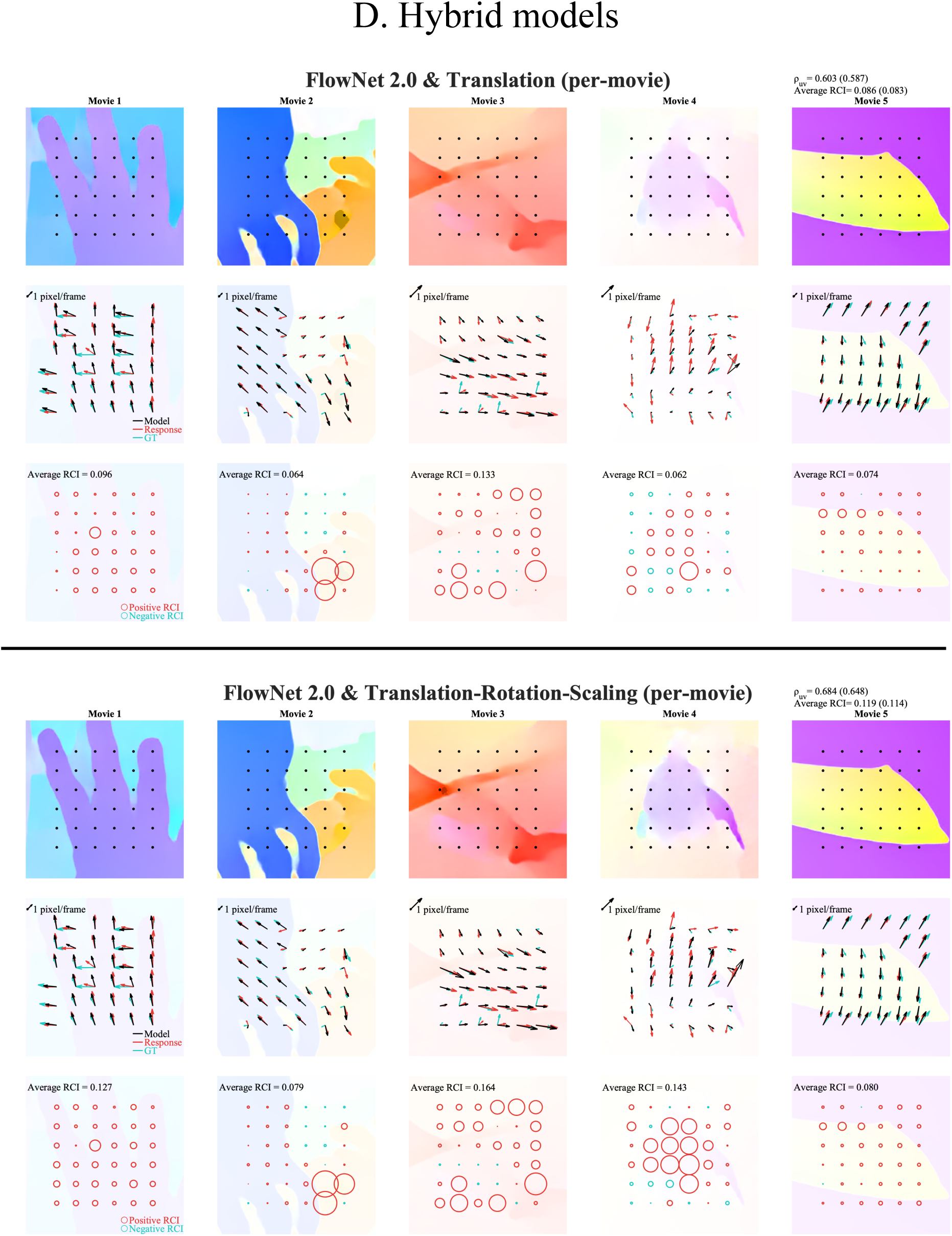
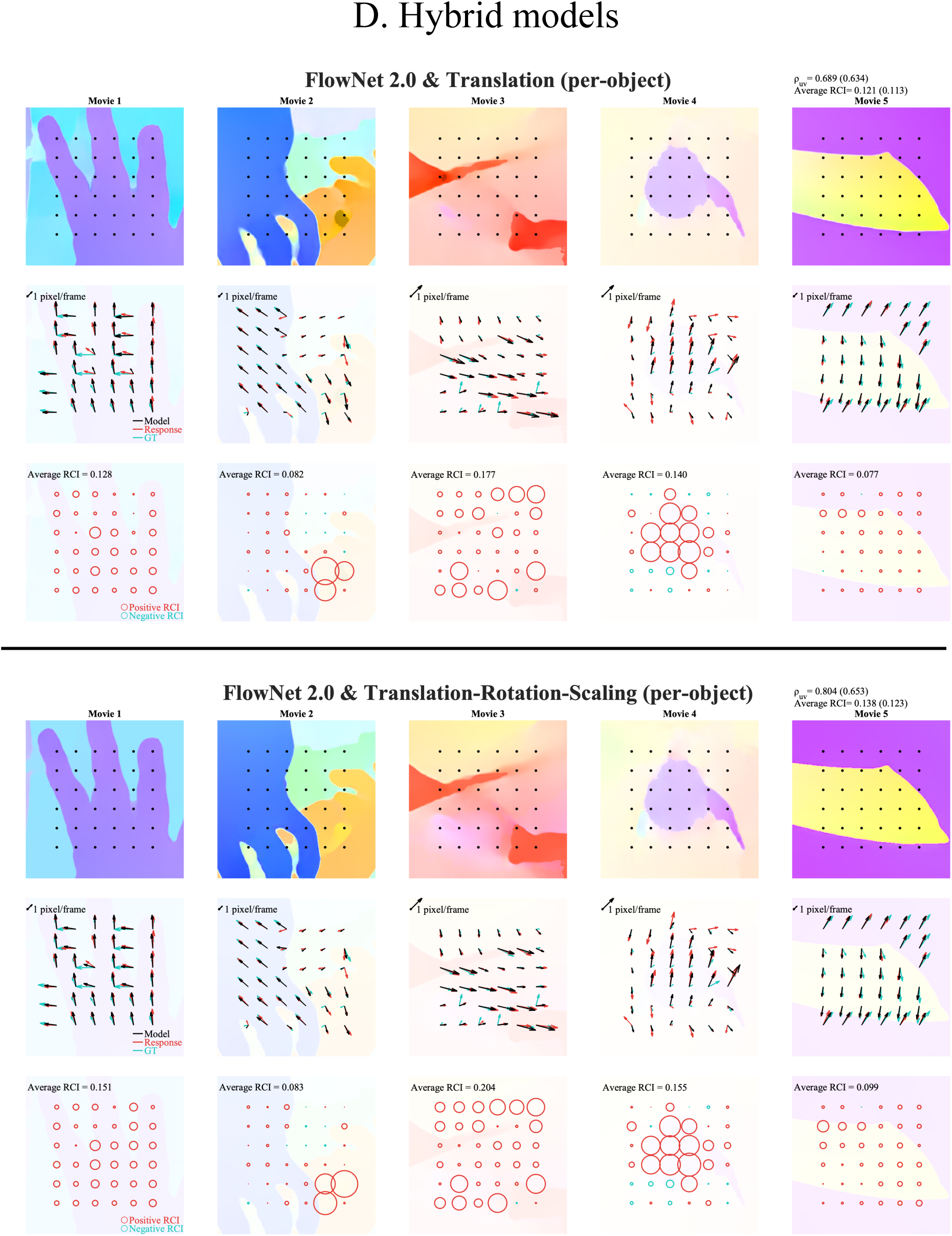
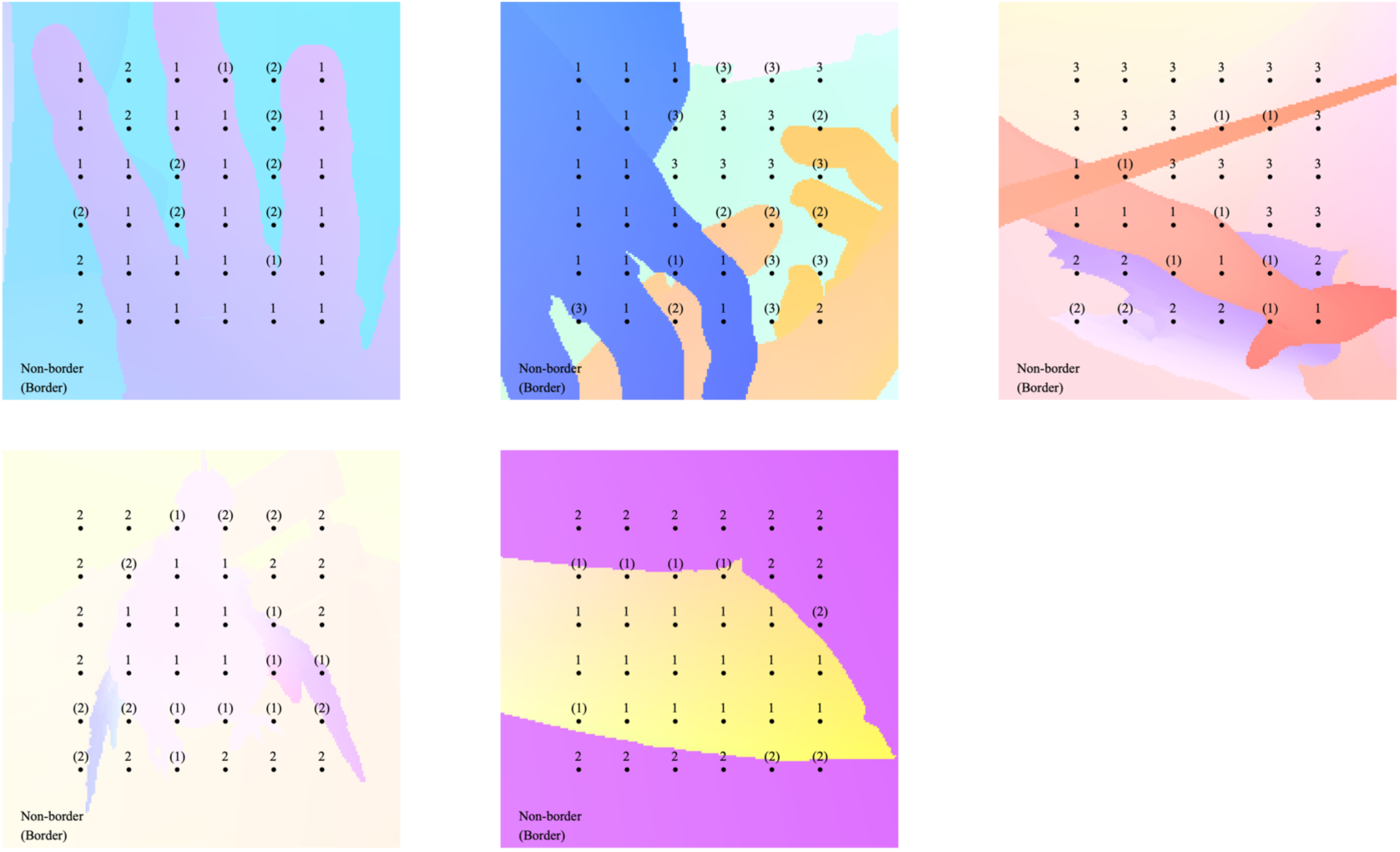
Classification of probed locations. The black dots represent probed locations. When data fitting to per-object transformation models, each location was classified as having one of two or three object layers, as indicated by the number. During evaluation of pooling models, locations near borders were considered to be border points, as indicated by the bracketed numbers.

**Table S1.** The Pearson correlation matrix and the partial correlation matrix between all pairs of four observers. The effects of GT are removed in the partial correlation.

| Correlation (Direction) |  |  |  |  |
| --- | --- | --- | --- | --- |
|  | Obs #1 | Obs #2 | Obs #3 | Obs #4 |
| Observer #1 |  |  |  |  |
| Observer #2 | 0.959 |  |  |  |
| Observer #3 | 0.958 | 0.941 |  |  |
| Observer #4 | 0.946 | 0.953 | 0.933 |  |
| The other Observers | 0.985 | 0.983 | 0.976 | 0.977 |
| Ground Truth | 0.955 | 0.945 | 0.942 | 0.962 |

| Correlation (Speed) |  |  |  |  |
| --- | --- | --- | --- | --- |
|  | Obs #1 | Obs #2 | Obs #3 | Obs #4 |
| Observer #1 |  |  |  |  |
| Observer #2 | 0.620 |  |  |  |
| Observer #3 | 0.653 | 0.695 |  |  |
| Observer #4 | 0.688 | 0.754 | 0.842 |  |
| The other Observers | 0.711 | 0.756 | 0.810 | 0.866 |
| Ground Truth | 0.784 | 0.737 | 0.791 | 0.868 |

| Correlation (uv components ) |  |  |  |  |
| --- | --- | --- | --- | --- |
|  | Obs #1 | Obs #2 | Obs #3 | Obs #4 |
| Observer #1 |  |  |  |  |
| Observer #2 | 0.805 |  |  |  |
| Observer #3 | 0.828 | 0.871 |  |  |
| Observer #4 | 0.816 | 0.872 | 0.891 |  |
| The other Observers | 0.852 | 0.897 | 0.915 | 0.913 |
| Ground Truth | 0.865 | 0.840 | 0.869 | 0.917 |

| Partial Correlation (Direction) |  |  |  |  |
| --- | --- | --- | --- | --- |
|  | Obs #1 | Obs #2 | Obs #3 | Obs #4 |
| Observer #1 |  |  |  |  |
| Observer #2 | 0.589 |  |  |  |
| Observer #3 | 0.590 | 0.461 |  |  |
| Observer #4 | 0.339 | 0.490 | 0.301 |  |
| The other Observers | 0.816 | 0.837 | 0.768 | 0.665 |

| Partial Correlation (Speed) |  |  |  |  |
| --- | --- | --- | --- | --- |
|  | Obs #1 | Obs #2 | Obs #3 | Obs #4 |
| Observer #1 |  |  |  |  |
| Observer #2 | 0.101 |  |  |  |
| Observer #3 | 0.086 | 0.269 |  |  |
| Observer #4 | 0.024 | 0.341 | 0.512 |  |
| The other Observers | 0.099 | 0.318 | 0.374 | 0.432 |

| Partial Correlation (uv components ) |  |  |  |  |
| --- | --- | --- | --- | --- |
|  | Obs #1 | Obs #2 | Obs #3 | Obs #4 |
| Observer #1 |  |  |  |  |
| Observer #2 | 0.291 |  |  |  |
| Observer #3 | 0.311 | 0.526 |  |  |
| Observer #4 | 0.115 | 0.472 | 0.477 |  |
| The other Observers | 0.317 | 0.583 | 0.594 | 0.481 |

**Table S2.**
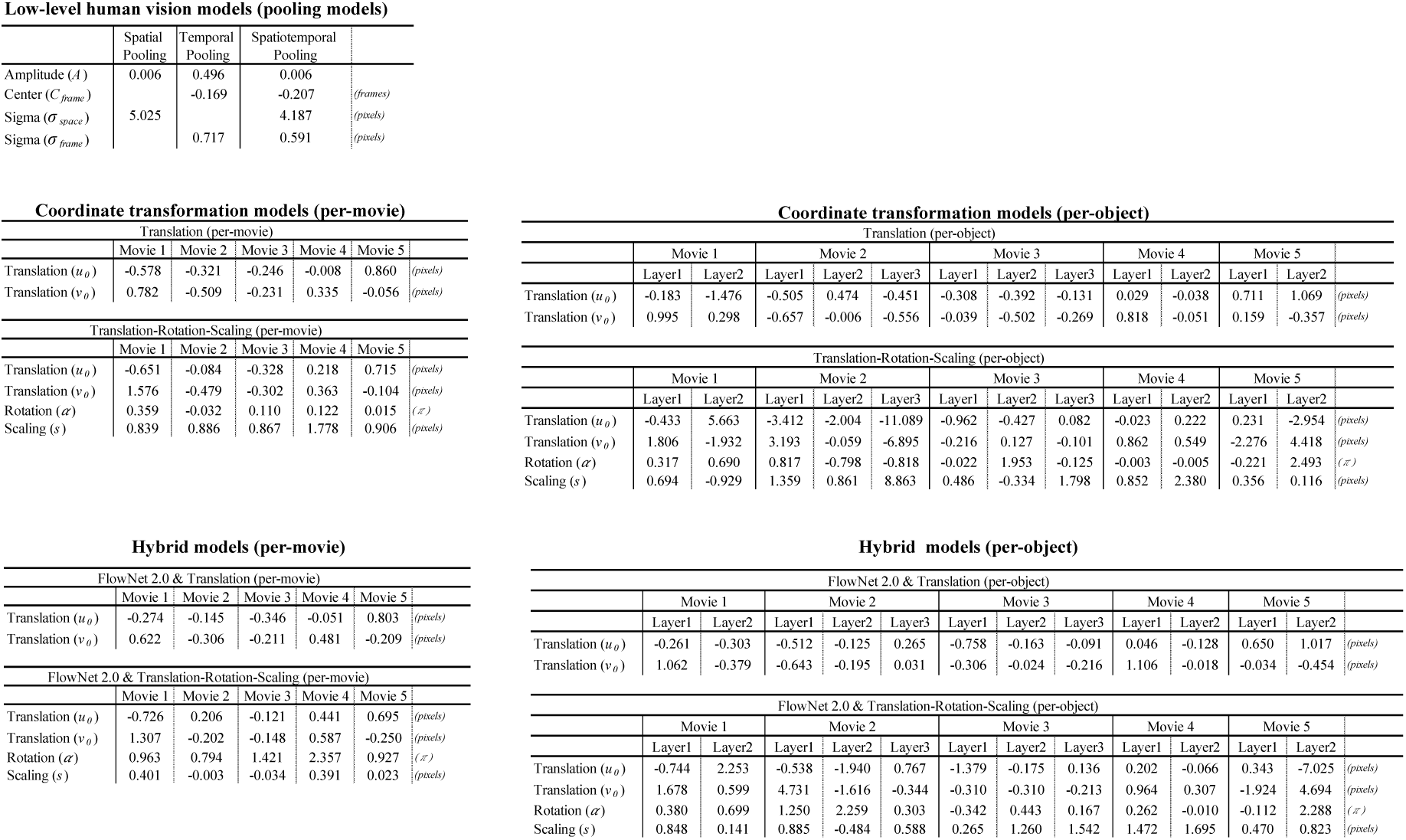
The best-fit parameters of all fitting models. All data points were used for fitting.

**Table S3.**
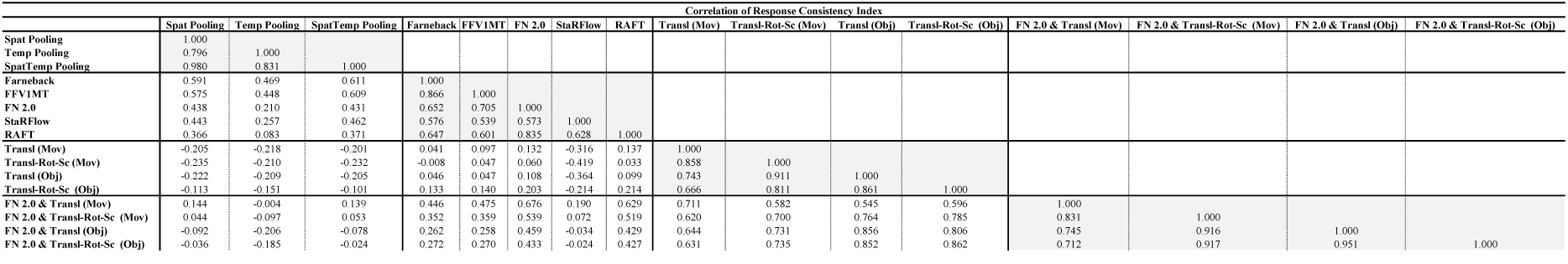
The correlations of the RCIs among models. We used 180 points (five movies x 36 locations) to calculate Pearson correlations between pairs of models. Spat: Spatial. Temp: Temporal. SpatTemp: Spatiotemporal. Transl: Translation. Rot-Sc: Rotation & Scaling. Transl-Rot-Sc: Translation, Rotation & Scaling. GL: Global. Obj: Per-Object.

**Table S4.** Performance index values of the spatial pooling models for non-border and border points, in the same format as Table 1.

|  | Boundary (Spatial Pooling) |  |  |  |  |  |  |  |  |  |  |  |
| --- | --- | --- | --- | --- | --- | --- | --- | --- | --- | --- | --- | --- |
| | Average EPE (GT) | Average EPE (Resp) | $r_{Dir}$ (GT) | $r_{Dir}$ (Resp) | $\rho_{Dir}$ | $r_{Spd}$ (GT) | $r_{Spd}$ (Resp) | $\rho_{Spd}$ | $r_{uv}$ (GT) | $r_{uv}$ (Resp) | $\rho_{uv}$ | Average RCI |
| Non-Border | 0.250 (0.247) | 0.809 (0.814) | 1.000 (1.000) | 0.959 (0.979) | 0.071 (0.063) | 1.000 (0.999) | 0.854 (0.924) | -0.106 (-0.061) | 0.959 (1.000) | 0.855 (0.948) | -0.023 (-0.039) | 0.004 (0.004) |
| Border | 0.553 (0.559) | 1.002 (1.020) | 0.965 (0.983) | 0.891 (0.943) | 0.330 (0.318) | 0.939 (0.966) | 0.663 (0.806) | 0.153 (0.111) | 0.882 (0.973) | 0.668 (0.862) | 0.417 (0.350) | 0.042 (0.041) |

## Notes

https://doi.org/10.17605/OSF.IO/BU7PD

### Competing Interest Statement

The authors have declared no competing interest.

